# Alternative splicing impacts microRNA regulation within coding regions

**DOI:** 10.1101/2023.04.20.536398

**Authors:** Lena Maria Hackl, Amit Fenn, Zakaria Louadi, Jan Baumbach, Tim Kacprowski, Markus List, Olga Tsoy

## Abstract

MicroRNAs (miRNAs) are small non-coding RNA molecules that bind to target sites in different gene regions and regulate post-transcriptional gene expression. Approximately 95% of human multi-exon genes can be spliced alternatively, which enables the production of functionally diverse transcripts and proteins from a single gene. Through alternative splicing, transcripts might lose the exon with the miRNA target site and become unresponsive to miRNA regulation. To check this hypothesis, we studied the role of miRNA target sites in both coding and noncoding regions using six cancer data sets from The Cancer Genome Atlas (TCGA). First, we predicted miRNA target sites on mRNAs from their sequence using TarPmiR. To check whether alternative splicing interferes with this regulation, we trained linear regression models to predict miRNA expression from transcript expression. Using nested models, we compared the predictive power of transcripts with miRNA target sites in the coding regions to that of transcripts without target sites. Models containing transcripts with target sites perform significantly better. We conclude that alternative splicing does interfere with miRNA regulation by skipping exons with miRNA target sites within the coding region.

## INTRODUCTION

MicroRNAs (miRNAs) are short (16-27 nucleotides (1)) non-coding RNAs that regulate post-transcriptional gene expression. They usually repress the target gene by destabilizing its transcript and/or by repressing its translation (2). Through a complementary target site (position 2-8 from the 5’ end, commonly referred to as seed sequence) they bind to their target mRNA and guide the RNA-induced silencing complex (RISC) to degrade it (3). In mammals, miRNAs regulate more than 60% of all protein-coding genes (4). They play an important role in health and disease. For example, tissue-specific miRNAs control cell differentiation (5) and miRNA downregulation is associated with tumorigenesis, e.g., the downregulation of liver-specific miRNA miR-122 in hepatocellular carcinoma (HCC) (6, 7). By analyzing the expression of 11 major human cancers from the Cancer Genome Atlas (TCGA), Li *et al*. (8) showed that the correlation between miRNA and target gene expression is reduced in tumors compared to normal tissue. Since individual miRNAs are able to simultaneously downregulate several target genes and thereby affect whole pathways, they are interesting therapeutical targets (9).

MiRNAs are known to bind to the 3’ untranslated region (3’-UTR) of their targets (10). However, Lytle *et al*. (11) moved a target site of let-7a miRNA from the 3’-UTR to the 5’-UTR in human HeLa cells and demonstrated that both 5’-UTR and 3’-UTR can be targeted. Lee *et al*. (12) found that not only the 5’-end of miRNAs can interact with the 3’-UTR of mRNAs but also *vice versa*. They identified many mRNAs that simultaneously contain 5’-end and 3’-end target sites enabling combinatorial interactions between a single miRNA and both UTRs of an mRNA. While previous experiments found the reduction of protein levels by around 40%–60% when using only 3’-UTR (5), the authors observed an even greater reduction of protein abundance by also including miRNA target sites in the 5’-UTR. They validated their findings experimentally using hsa-miR-34a binding to both 3’-UTR and 5’-UTR of AXIN2.

A gene’s coding region can also contain potential miRNA target sites. Forman *et al*. (13) analyzed publicly available proteomics datasets and demonstrated that miRNA target sites in coding regions are functional but less conserved and effective in repression or inhibition of target genes than 3’-UTR sites. Hausser *et al*. (14) analyzed putative miRNA target sites in coding regions that were predicted computationally or inferred based on expression changes upon miRNA transfection. Target sites in the coding regions were found to have a smaller impact on mRNA stability but to be more effective in inhibiting translation, while 3’-UTR sites trigger mRNA degradation more efficiently. The authors concluded that a combination of both enables fine-tuning of the miRNA regulatory effects.

The miRNA-mediated regulation through interaction with target sites in the coding regions might be affected by alternative splicing. Approximately 95% of human multi-exon genes can be spliced alternatively (15), which leads to different combinations of exons in resulting transcripts. If the exon with the miRNA target site is spliced out, the transcript might evade miRNA regulation. However, the impact of alternative splicing on miRNA-mediated mRNA regulation has been previously addressed only in one study. Han *et al*.

(16) studied the impact of alternative splicing and alternative polyadenylation of 3’-UTRs on miRNA-mediated repression efficiency in bladder cancer. They demonstrated that miRNA might fail to regulate alternatively spliced transcripts missing 3’-UTR exons.

The following open questions remain: Does alternative splicing of exons in the coding region affect miRNA-mediated regulation? How does this effect generalize to other tissues and conditions?

We address these questions and evaluate the impact of alternative splicing of coding regions on miRNA regulation on a transcriptome-wide scale for several types of cancer. To that end, we computationally predict miRNA target sites using TarPmiR (17) and filter out miRNA-gene pairs based on their expression level and correlation between gene and miRNA expression. We then construct nested linear regression models to predict miRNA expression based on the expression of transcripts with and without miRNA target sites and conclude that alternative splicing does indeed interfere with miRNA-mediated mRNA regulation.

## MATERIALS AND METHODS

### Workflow

Figure 1 illustrates the workflow we used to investigate the impact of alternative splicing on miRNA-mediated gene expression regulation. Human miRNA and mRNA sequences were input to TarPmiR for miRNA target site prediction on the mRNAs (Figure 1.1, for details see Methods below). Based on the binding probability and the location of a target site for each miRNA-transcript pair, we categorized transcripts into four types (Figures 1.2, 2A): a) non-binding transcripts;

- transcripts with binding in the coding region; c) transcripts with binding in non-coding regions; d) transcripts with binding in both coding and non-coding regions. The number of miRNA-transcript pairs in each category is shown in Figure 3B. To investigate the role of miRNA target sites in coding regions opposite to non-coding regions, from here on the same steps were performed separately for each of the three settings (Figure 2B):
- all transcripts (ALLT): the full models contain all transcripts; the reduced models contain non-binding transcripts. This setting allows us to investigate the role of miRNA target sites independent of their location in the gene.
- transcripts not binding in non-coding regions (TNBN): the full models contain non-binding transcripts and transcripts only binding in coding regions; the reduced models contain non-binding transcripts. This setting allows us to investigate the impact of miRNA target sites in coding regions in the absence of miRNA target sites in non-coding regions.
- transcripts binding in non-coding regions (TBN): the full models contain transcripts binding in non-coding regions and transcripts with binding in both coding and non-coding regions; the reduced models contain transcripts binding in non-coding regions. This setting allows us to investigate the impact of miRNA target sites in coding regions in the presence of miRNA target sites in the non-coding regions.

**Figure 1.**
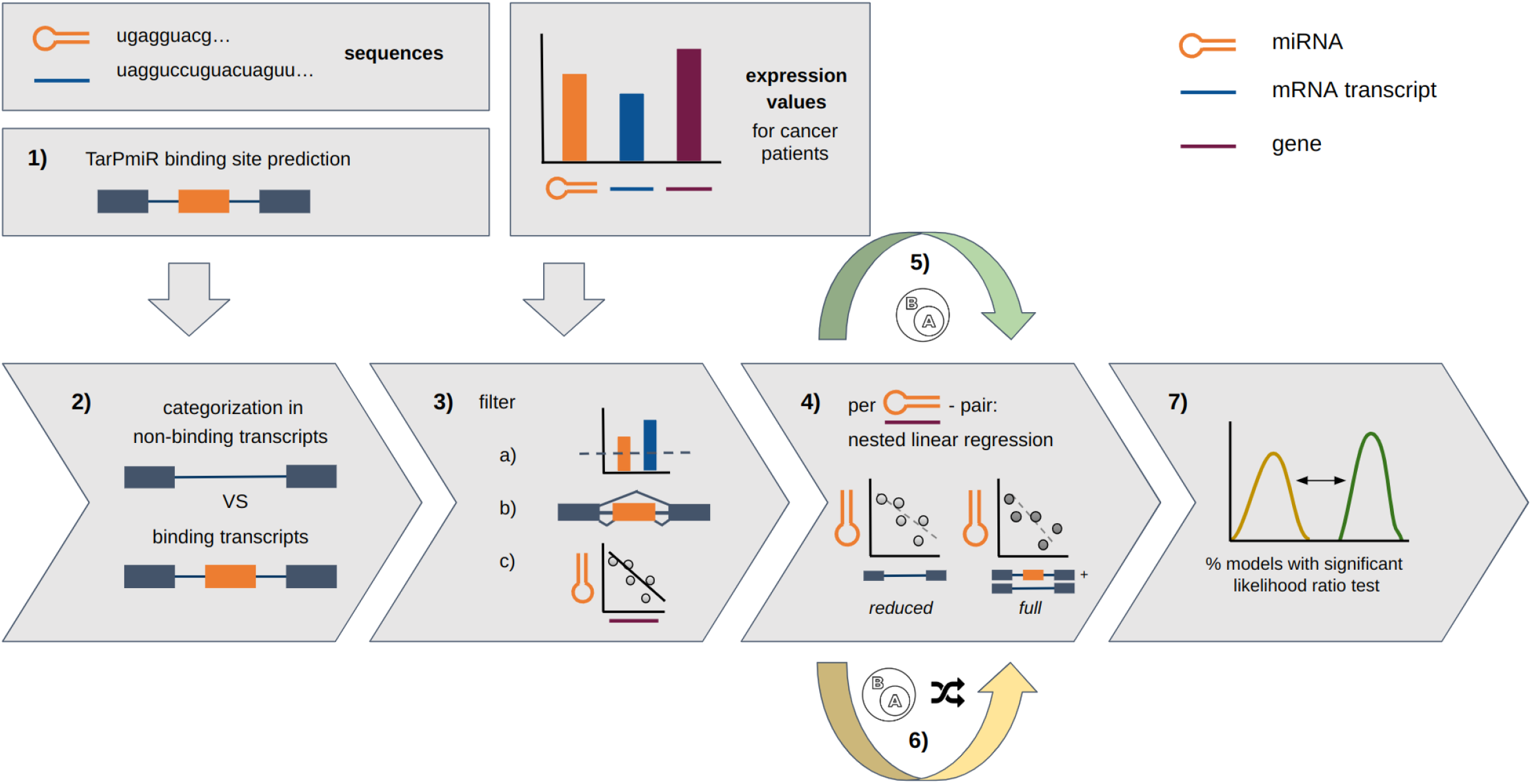
Analysis pipeline using miRNA and mRNA expression and sequence data. 1) TarPmiR target site prediction, 2) categorization in non-binding *vs*. binding transcripts, 3) filtering a) for expression and variance above the chosen thresholds (see Methods), b) alternatively spliced genes, c) negative correlation of miRNA and gene expression, 4) per miRNA-gene pair nested linear regression: non-binding transcript regression and all transcript regression, 5) subsampling of nested models, 6) subsampling and label randomization of nested models, 7) likelihood ratio test between nested model pairs. The pipeline was run for the three settings ALLT (all transcripts), TNBN (transcripts not binding in non-coding region) and TBN (transcripts binding in non-coding region) separately from step 2) on.

**Figure 2.**
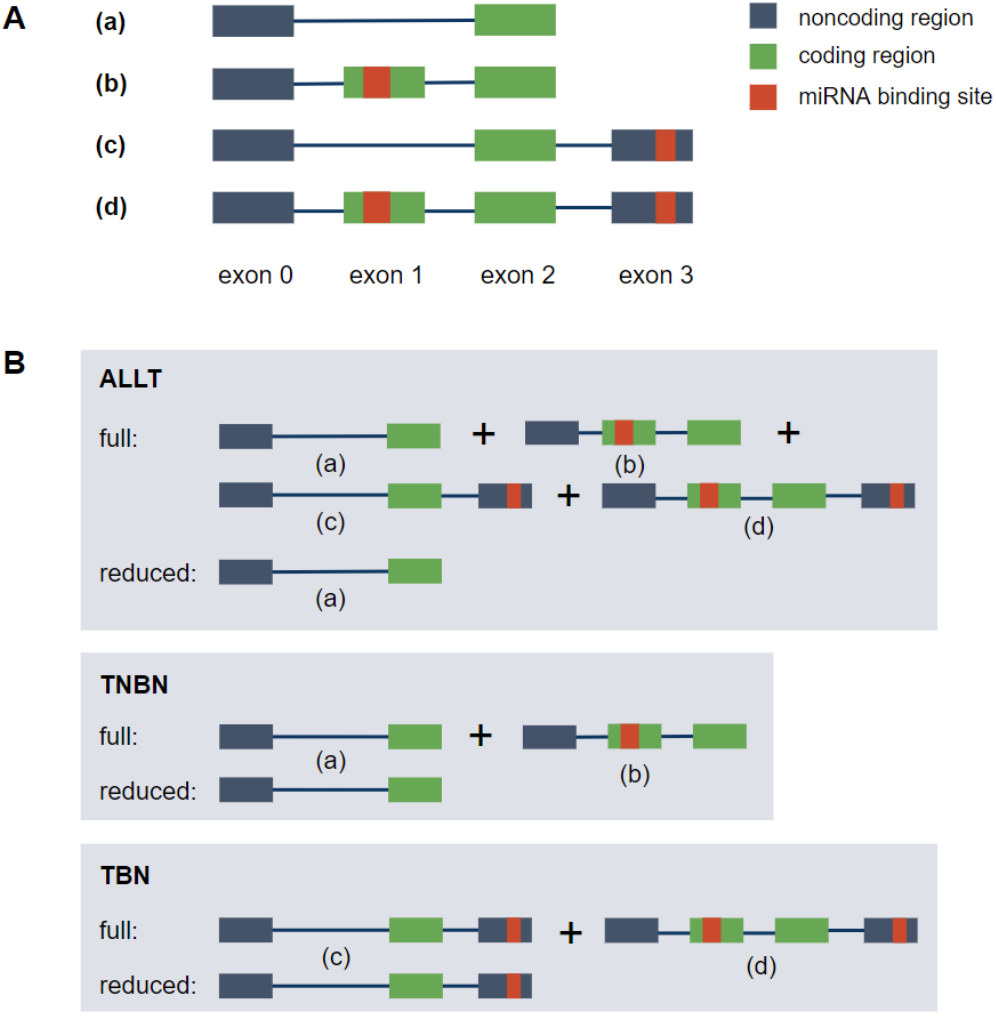
**A** Transcripts are divided into four different transcript types: (a) non-binding transcripts, (b) transcripts with target sites only in the coding region, (c) transcripts with target sites only in the non-coding region (3’ -UTR or 5’ -UTR), (d) transcripts with target sites in both the coding and non-coding region **B** Structure of the nested miRNA-gene-level linear regression models. The full model is trained on: all transcripts (ALLT), only transcripts without target sites in non-coding region (TNBN), and only transcripts with target sites in non-coding region (TBN). Accordingly, the reduced model is only trained on a subset of the transcripts without target sites in the investigated region.

**Figure 3.**
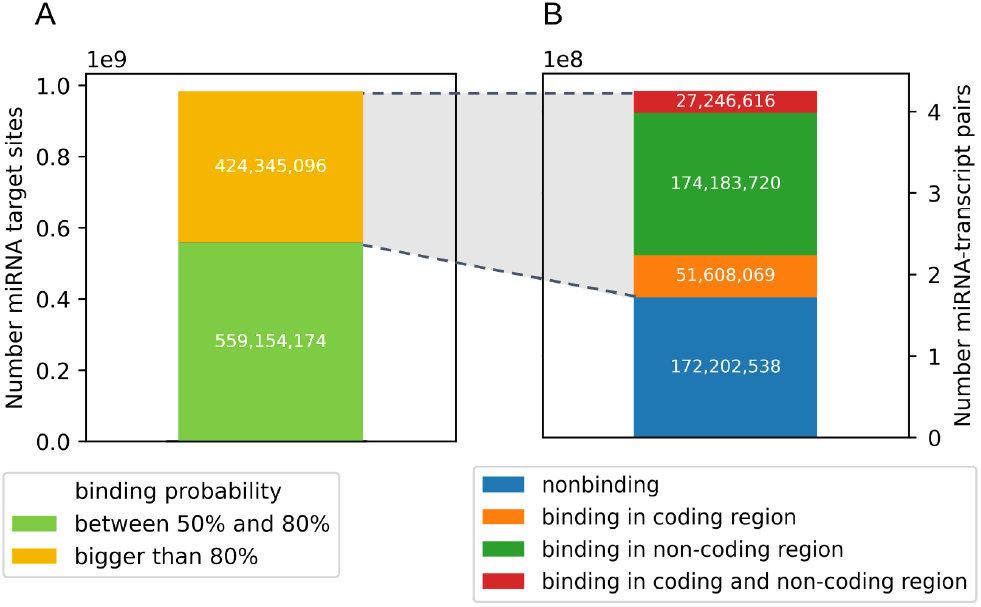
Categorization of predicted TarPmiR target sites before any filtering **A** in target sites with binding probability between 50% and 80% and above 80% and **B** in miRNA-transcript pairs based on target region (coding/non-coding region).

After filtering (Figure 1.3), we constructed nested linear regression models for each remaining miRNA-gene pair to predict miRNA expression from transcript expression (Figure 1.4) and filtered the models by the root mean squared error (RMSE) (Figure S2). This procedure was repeated on both random subsets (Figure 1.5) and random subsets with additionally permuted labels (Figure 1.6). Finally, between each pair of full and reduced models we performed a likelihood ratio test (Figure 1.7).

**Figure 4.**
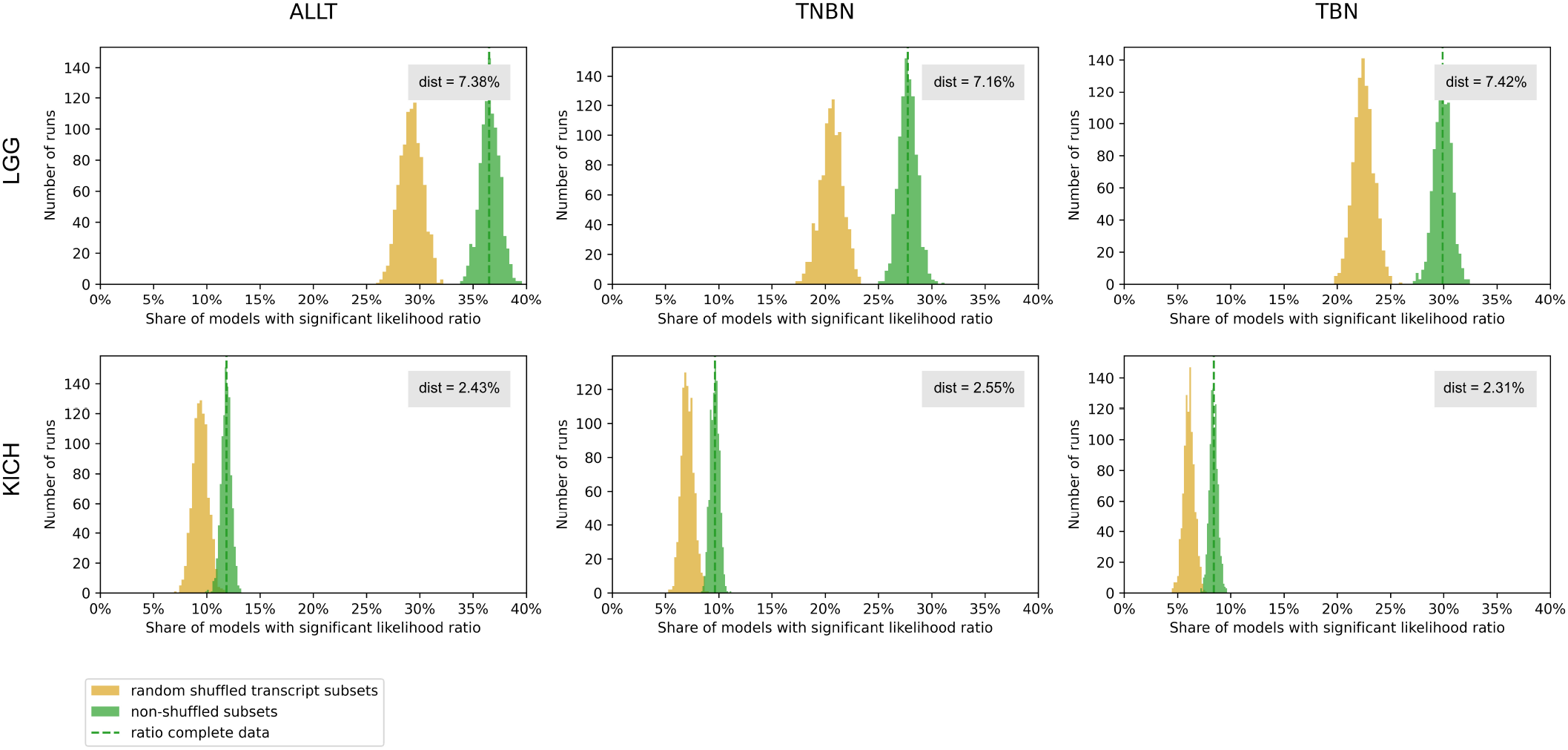
The ratio of models with statistically significant (<0.05) corrected p-values of the likelihood ratio test statistic calculated between nested regression models is shown as the dashed green line. To estimate the distribution, the ratio was calculated 1,000 times for random subsets of miRNA-gene pairs (green histogram) and to estimate the impact of alternative splicing, the ratio was calculated 1,000 times for random subsets of miRNA-gene pairs while randomizing the transcript binding labels within a gene (yellow histogram). Dist describes the difference between the average ratio of models based on subsampled real miRNA-gene pairs and after randomizing the transcript category labels. This is shown separately for diseases Brain lower grade glioma (LGG) and Kidney chromophobe carcinoma (KICH) for settings ALLT, TNBN, and TBN.

### Prediction of miRNA target sites

#### Sequence data

We downloaded 2,656 human mature miRNA sequences from miRBase (18) (release 22.1). The Ensembl database (19) release 100 (GRCh38.p13 assembly) was used as a source for the 249,750 mRNA sequences (228,116 primary assembly sequences + 21,634 alternative assembly sequences), as well as coding region annotation. We converted all uracil bases to thymine bases as cDNA format is necessary for miRNA target site prediction.

#### miRNA target site prediction with TarPmiR

For prediction of miRNA target sites, we used the state-of-the-art tool TarPmiR (17). We chose TarPmiR because of its better recall and precision compared to other commonly used methods such as TargetScan (20), miRanda (21), and miRmap (22). We executed TarPmiR on the Ensembl mRNA sequences and miRBase mature miRNA sequences using the default parameters. The final output file contains miRNA target site candidates with binding probabilities > 50% (Figure S1). As computational prediction of target sites can lead to false-positive predictions, we kept only transcripts that contain target sites with binding probabilities > 80% (‘binding transcripts’) and transcripts that contain no target sites with binding probabilities > 50% (‘non-binding’ transcripts) (Figure 3A). The TarPmiR default is 50% but we raised the threshold to 80% to focus on high-confidence predictions (Figure S3). Transcripts that contain only target sites with a binding probability between 50% and 80% were filtered out to avoid noise.

The predicted target sites were then mapped back onto the exons and categorized into target sites in the coding region and target sites in the non-coding region (5’-UTR and 3’-UTR). Target sites overlapping both coding and non-coding regions were assigned to both categories.

#### Expression filter

To further reduce the number of potential false-positive predictions, we applied an expression filter. MiRNA and mRNA expression data were collected from The Cancer Genome Atlas (TCGA). We used the Xena platform (23) to download the batch effect corrected, TPM normalized, and log-transformed gene expression data (version 2016-09-01), transcript expression data (version 2019-02-25) and miRNA mature strand expression data (version 2016-12-29) from the TCGA Pan-Cancer (PANCAN) cohort. We investigated the following cancer types: Brain lower grade glioma (LGG), Kidney chromophobe carcinoma (KICH), Liver hepatocellular carcinoma (LIHC), Kidney renal cell carcinoma (KIRC), and Breast Invasive Carcinoma: Invasive Lobular Carcinoma (ILC) and Invasive Ductal Carcinoma (IDC). We selected tissues with a high proportion of alternatively spliced genes such as Brain tissue (LGG), Liver tissue (LIHC) and Kidney tissue (KICH, KIRC). Breast tissue (IDC, ILC) was chosen for comparison due to the high number of samples available in TCGA (24). We filtered out miRNAs with expression variance smaller than 0.2 between samples within a cancer type dataset to reduce noise and to prevent overfitting of the model. We filtered out genes and transcripts that are not expressed in 25% or more samples within a dataset.

#### Alternative splicing filter

To account for alternative splicing, we kept only miRNA-gene pairs where a gene has at least one transcript containing a miRNA target site with TarPmiR probability > 80% in the investigated region and at least one other transcript containing no miRNA target site with TarPmiR probability > 50% in the same investigated region.

#### Correlation filter

miRNAs most frequently repress the expression of their targets (25, 26). Therefore, we expect a negative correlation between miRNA expression and target gene expression. To focus on the down-regulating effect that most miRNAs have on target gene expression, the Pearson standard correlation coefficient between miRNA expression and gene expression was calculated for all miRNA-gene pairs. We kept only pairs with a negative Pearson correlation coefficient for further analysis.

### Nested linear regression on expression

#### Full and reduced models

The samples for all miRNA-gene pairs were divided into training (80%) and test sets (20%). We constructed nested linear regression models to predict miRNA expression from transcript expression. Per miRNA-gene pair a full model was trained on all transcripts and accordingly a reduced model was trained on the transcripts without target sites in the investigated region, both using the ordinary least squares method (see Workflow Description below).

For the resulting models, the RMSE was calculated on the test sets. All nested models with an error smaller than 0.7 for the reduced and corresponding full models were kept. The threshold was set by visually examining the number of models left after filtering (Figure S2). The likelihood ratio test was used between the reduced and full model, applying the Benjamini Hochberg multiple testing correction with a family-wise error rate of a= 0.05. We then calculated the ratio of nested models with statistically significant adjusted p-values (p<0.05), meaning nested models, where the full model outperformed the reduced model.

To estimate the distribution of ratios for the data set, we randomly sampled 10,000 miRNA-gene pairs 1,000 times and calculated the ratio of nested models, where the full model outperformed the reduced model.

#### Randomization

To show that the obtained ratio of nested models with significant adjusted p-value (p<0.05) cannot be reached by chance, we repeated the subsampling procedure described above with 10,000 miRNA-gene pairs 1,000 times, but shuffled the transcript binding labels within a gene while preserving the absolute number of binding transcripts and the absolute number of non-binding transcripts per gene. Per iteration, the reduced models were retrained and filtered by RMSE, as described above. For the remaining nested models, the likelihood ratio test statistic was calculated. We obtained the ratio of nested models with significant p-values after Benjamini Hochberg multiple testing correction as for the non-randomized models above.

## RESULTS

### Computational prediction of miRNA-gene pairs

Starting from 2,656 miRNA and 249,750 mRNA sequences TarPmiR predicted 983,499,270 target sites with probability > 50% (see Figure Sl), among those 424,345,096 target sites with probability> 80%. In total this amounted to 288,056,794 miRNA-transcript pairs with target sites. This high number might indicate a high number of false-positive predictions (Figure S4). In particular, target site predictions are sequence specific and hence do not account for a lack of expression of either a miRNA or its target in a specific condition, cell type or tissue. Thus, we used miRNA and mRNA expression data for the six cancer types from TCGA.

The expression matrices contain 178,927 transcripts of 57,471 genes before the expression filter. Table 1 shows the number of samples per cancer type dataset with both miRNA and gene expression. The number of miRNA samples after filtering by expression variance slightly differs between datasets (Table 1). Table 1 illustrates the number of miRNA-gene pairs after filtering, while Table Sl shows the intermediate numbers of miRNA-gene pairs after the single filtering steps and Figure S5 additionally shows the distribution of the number miRNAs per gene and *vice versa* after filtering. We filtered for miRNA-gene pairs with expression above the chosen thresholds, alternative splicing, and negative Pearson correlation between miRNA and gene expression (see Methods).

**Table 1.** Number of samples, miRNAs, and miRNA-gene pairs after all filtering steps shown for settings ALLT, TNBN, TBN and for the investigated cancer types.

| Cancer | # of samples | # of miRNAs | # of miRNA-gene pairs after all filters |  |  |
| --- | --- | --- | --- | --- | --- |
|  |  |  | ALLT | TNBN | TBN |
| LGG | 517 | 541 | 2,116,741 | 824,021 | 482,012 |
| KICH | 89 | 479 | 1,252,924 | 476,494 | 300,592 |
| LIHC | 413 | 623 | 1,411,890 | 515,100 | 321,083 |
| KIRC | 565 | 546 | 1,809,116 | 678,231 | 409,757 |
| ILC | 179 | 544 | 2,092,222 | 802,642 | 487,659 |
| IDC | 628 | 602 | 2,168,203 | 837,560 | 509,879 |

### Impact of alternative splicing on miRNA-mediated regulation

Previous studies focused on the role of miRNA target sites in non-coding regions (10, 11, 12). First, using the ALLT setting, we investigated the impact of miRNA binding on transcript expression independent of the location of target sites. This setting might reflect the mixed effect, where target sites in the non-coding region have more impact compared to target sites in the coding regions. To check this hypothesis, we developed two other settings described below.

Target sites in the coding regions might be spliced out due to alternative splicing, and if those target sites are important for miRNA-mediated regulation, the resulting transcript could evade miRNA regulation, i.e., its expression will be independent of miRNA regulation. To investigate this possibility, we used the TNBN setting.

Transcripts often contain a mixture of miRNA target sites-both in coding and non-coding regions. Target sites in the non-coding regions might fully overpower the effect of target sites in coding regions. To investigate this possibility, we used the TBN setting.

We calculated the likelihood ratio test between the nested models for each miRNA-gene pair and compared the ratio of models, where the full model statistically significantly outperformed the reduced model for ALLT, TNBN, and TBN, respectively. Next, we compared this ratio with the ratio of such models after randomizing transcript category labels (Figure 4 for LGG and KICH, Figure S6 for LIHC, KIRC, ILC, IDC).

The full models were found to consistently outperform the reduced models, whereas the magnitude of performance gain varies between cancer types. For Invasive Lobular Carcinoma (ILC) the effect is the weakest and the distribution of the ratios overlaps between real and randomized experiments. In the setting TNBN we see that also target sites in the coding region have an effect, supporting the notion that active miRNA target sites are not only found in non-coding regions.

For all analyzed cancer types and settings (ALLT, TNBN, TBN), this difference is significant (Mann-Whitney U test p-value < 0.05). This observation supports the hypothesis that on transcriptome-wide scale, alternative splicing impacts miRNA regulation by splicing out miRNA target sites in the coding regions.

### Gene Set Enrichment analysis

We performed a Gene Set Enrichment analysis of the top 500 genes that predict miRNA expression with most significant p-values using the Molecular Signatures Database (27) (28) for all six cancer types and settings. We overlayed the top 500 genes with all non-computational collections (C1, C2, C3, C5, C6, C7 (29), C8, H (30)). For all the cancer types and settings, we found cancer-related functions within the top ten most significantly enriched genesets of the non-computational collections for all diseases and settings besides for LGG setting TNBN (Supplementary Tables S2 - S19). Of those, we found an overlap with a geneset related to the specific cancer type for cancer type IDC in all settings, for ILC settings TBN and ALLT, for KIRC setting ALLT, and for LIHC setting TBN.

While overlaying the top 500 genes of all the cancer types and settings with only oncogenic genesets (C6), we found significant overlaps for all cancer types and settings (Supplementary Tables S20 - S37).

## DISCUSSION

We investigated the impact of miRNA target sites in the coding regions based on miRNA and mRNA expression data using a nested linear regression approach. On a transcriptome-wide scale, we demonstrated that alternative splicing impacts miRNA regulation, putatively by splicing out miRNA target sites in the coding regions. We observe the same effect in six cancer types.

While cancer-related terms were frequently observed among the top ten genesets in the Gene Set Enrichment analysis, several other enriched terms lacked a direct association with cancer. These likely indicate tissue-specific effects of miRNA regulation, which suggests that the interplay of miRNA regulation and AS probably also plays a role in tissue differentiation (31).

The interplay between miRNA regulation and alternative splicing in cancer development has rarely been addressed (16). However, several miRNA-gene pairs, for which we showed a significant impact of alternative splicing, have been demonstrated to be important for cancer subtypes.

It is known that microRNA-200 family miRNAs target genes ZEB1 and ZEB2, which both are involved in EMT and tumour metastasis (32). In our analysis, we found that miRNA miR-141-5p regulates ZEB2 (p&x2248; 2.39e−4 in LGG setting TBN and p&x2248; 9.00e−5 in LGG setting ALLT). miR-141-5p inhibits glioma cell growth and migration by repressing ZEB1 expression (33). In pancreatic cancer, however, treatment of MiaPaCa-2 cells with gemcitabine caused an upregulation of the ZEB1 protein through alternative polyadenylation of the transcript (34). Thereby the ZEB1 3’-UTR was shortened and miRNA target sites in the last exon deleted. We were able to observe gene ZEB1 evading regulation by miR-141-5p through alternative splicing (p&x2248; 3.32e−5 in LGG using setting TBN).

Tumor suppressor miR-30c is known to inhibit prostate cancer by targeting the 3’-UTR of the SRSF1 splicing factor oncoprotein to downregulate its expression in prostate cancer (35). This expression is correlated with the pathological stage of prostate cancer and biochemical recurrence. SRSF1 is also known to be over-expressed in kidney tumor (36). We found the miRNA miR-30c-1-3p and gene SRSF1 interaction significant in KICH setting TNBN (p&x2248; 2.39e−2) and in KIRC setting ALLT (p&x2248; 3.03e−2). In renal cancer 3’-UTR variants of SRSF1 were discovered with differing miRNA target sites (37), a differential regulation mechanism potentially existing for miR-30c as well. We found SRSF1 also interacts with miR-7 in lower grade glioma - more specifically with miR- 7-2-3p (p&x2248; 9.22e−8 for setting ALLT). The splicing factor SRSF1 transcript, besides being repressed by miR-7, is also targeting the miRNA through binding, thereby generating a negative feedback loop (38).

In KIRC setting ALLT we found miR-18a-3p regulating K-Ras expression (p&x2248; 1.41e−3), an interaction which was previously shown experimentally (39). MiR-18a* acts as a tumor suppressor by targeting oncogene K-Ras. K-Ras is known to be alternatively spliced into two isoforms K-Ras 4B, which is anti-apoptotic and ubiquitously expressed, and K-Ras 4A, which is pro-apoptotic and expressed in only a subset of tissues such as kidney, lung and colon (40). In renal cell carcinoma oncogene K-Ras 4A was observed as upregulated and the isoform’s influence on cell survival and proliferation shown (41).

Our study provides the first steps toward investigating the role of alternative splicing in miRNA gene regulation. In our analysis, we consider miRNA-gene interactions as binary, while genes acting as competing endogenous RNAs actually form a complex gene-regulatory network based on miRNA competition (42, 43). We chose a simple binary model over a more complex network model with n-to-n interactions as the results are easier to interpret and clearly support our findings. However, further work is needed to understand the impact of AS on miRNA regulation on a network level. Furthermore, many transcripts show target sites for several miRNAs (Figure S7). The multi-mRNA effect should be taken into account in the future to refine the miRNA-transcript relationship. In this study, we focused exclusively on exon sequences for miRNA target site prediction. Further work is needed to elucidate the role of alternative splicing events other than exon skipping, such as intron retention, on miRNA regulation. Another interesting aspect worth considering in the future is the question of the effect of miRNA regulation on the alternative splicing machinery, as miRNAs can bind to splicing factors and alter splicing activity (44). This work focuses on the more common miRNA-mediated downregulation rather than upregulation, as miRNA-mediated upregulation is rare and currently not sufficiently understood (25, 45, 46).

The current availability of miRNA and mRNA expression data from the same tissue and condition limits the study. However, the effect is clearly seen in different cancer types, which suggests that this effect might be commonly observed. The most promising miRNA-gene pairs might be feasible for experimental validation. However, the demonstrated effect has a transcriptome-wide scale meaning that might not be clearly observed for single miRNA-gene pairs. Nevertheless, we published the list of significant cancer-specific miRNA-gene interactions affected by alternative splicing (see Data Availability), and provide a basis for further experimental investigation of specific interactions and the influence of alternative splicing. In the future, we are planning to provide our findings as a user-friendly database where the miRNA-gene pairs can be investigated visually.

## CONCLUSION

We have developed a new computational method to assess the influence of miRNA binding in coding regions on the whole transcriptome. We studied the impact of alternative splicing on miRNA regulation on the whole transcriptome for several cancer types while focusing on the coding region. Using sequence data, miRNA target sites were predicted on human mRNA and the difference in correlation between miRNA expression and binding *vs*. non-binding transcript expression was investigated using nested linear regression models. We were able to show that miRNAs binding in coding regions are effective at reducing transcript expression and that transcripts that splice out the miRNA target site are less affected by miRNA-mediated downregulation. Beyond the influence of alternative splicing, we show evidence that the coding region plays a role in miRNA regulation. Our findings suggest that further clinical studies can be directed at studying miRNA target sites in the coding region.

## DATA AVAILABILITY

All data presented are derived from previously published data sets as indicated. The used input data and output data including miRNA-gene pairs with p-values and the results of the Gene Set Enrichment analysis for all six investigated cancer types are available at https://doi.org/10.6084/m9.figshare.21821181.v2. The code used for this study was written in Python and is available at https://github.com/CGAT-Group/miRNA-AS.

## ACKNOWLEDGEMENTS

The authors thank the anonymous reviewers for their valuable suggestions. This work was supported by the German Federal Ministry of Education and Research (BMBF) within the framework of the e:Med research and funding concept (grants 01ZX1908A and 01ZX2208A). This work was also developed as part of the ASPIRE project and is funded by the German Federal Ministry of Education and Research (BMBF) under grant number 031L0287B. J.B. was partially funded by his VILLUM Young Investigator Grant (nr. 13154). The results published here are in part based upon data generated by the TCGA Research Network: https://www.cancer.gov/tcga.

