## Supplementary Data and Figures for "Alternative splicing impacts microRNA regulation within coding regions"

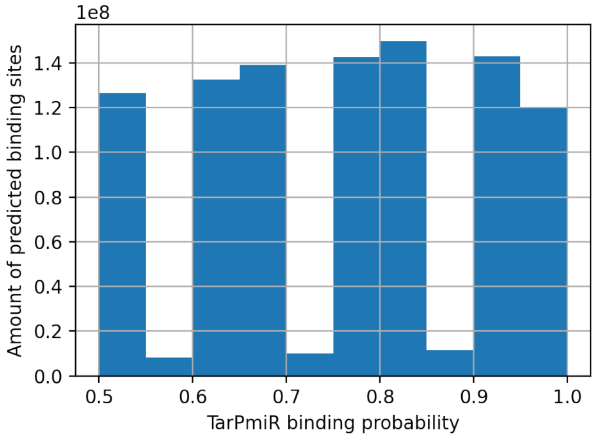


**Figure S1.** Binding probabilities of predicted TarPmiR target sites before any filtering

**Table S1.** Intermediate number of miRNA-gene pairs after different filtering steps and number final nested models shown for settings ALLT, TNBN, TBN and for the investigated cancer types.

| | # of miRNA-gene pairs after expression filter | | | # of miRNA-gene pairs after alternative splicing filter | | | # of miRNA-gene models after RMSE filter | | |
|---|---|---|---|---|---|---|---|---|---|
| Cancer | ALLT | TNBN | TBN | ALLT | TNBN | TBN | ALLT | TNBN | TBN |
| LGG | 5,537,553 | 2,580,282 | 1,416,480 | 3,926,467 | 1,507,316 | 913,827 | 610,263 | 237,734 | 153,394 |
| KICH | 4,818,224 | 2,268,596 | 1,261,454 | 3,252,566 | 1,244,890 | 765,695 | 546,157 | 213,742 | 147,345 |
| LIHC | 5,970,450 | 2,824,880 | 1,557,801 | 3,347,063 | 1,217,406 | 759,370 | 313,382 | 114,776 | 83,152 |
| KIRC | 5,534,423 | 2,586,037 | 1,428,360 | 3,849,318 | 1,470,433 | 903,506 | 704,955 | 263,378 | 170,929 |
| ILC | 5,527,849 | 2,592,399 | 1,435,927 | 3,849,351 | 1,478,371 | 908,345 | 715,294 | 282,035 | 190,513 |
| IDC | 6,057,604 | 2,841,251 | 1,575,977 | 4,115,021 | 1,578,926 | 967,695 | 469,066 | 181,298 | 108,074 |

*2*

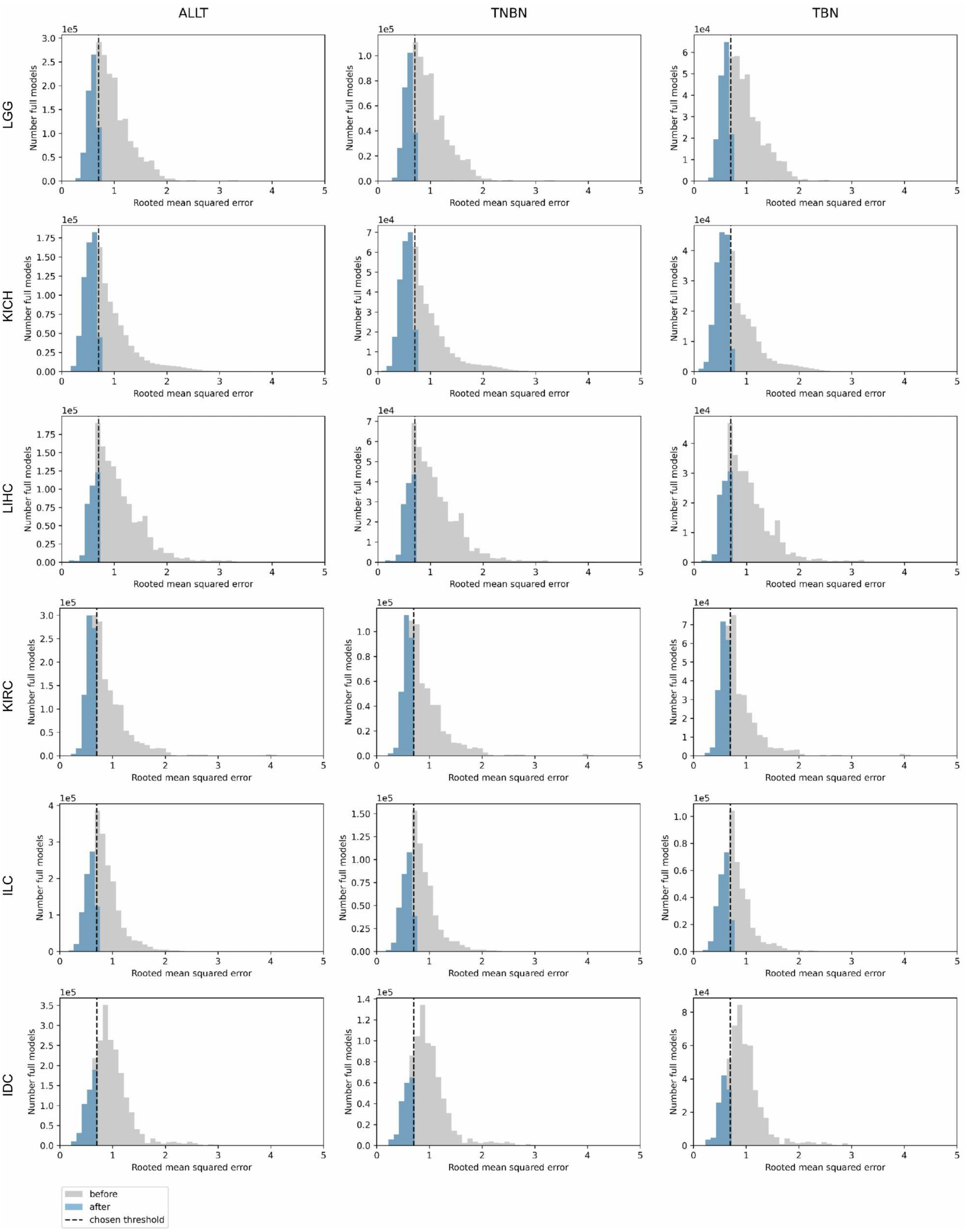

ALLT
TNBN
TBN
LGG
KICH
LIHC
KIRC
ILC
IDC
Number full models
Rooted mean squared error
before
after
chosen threshold

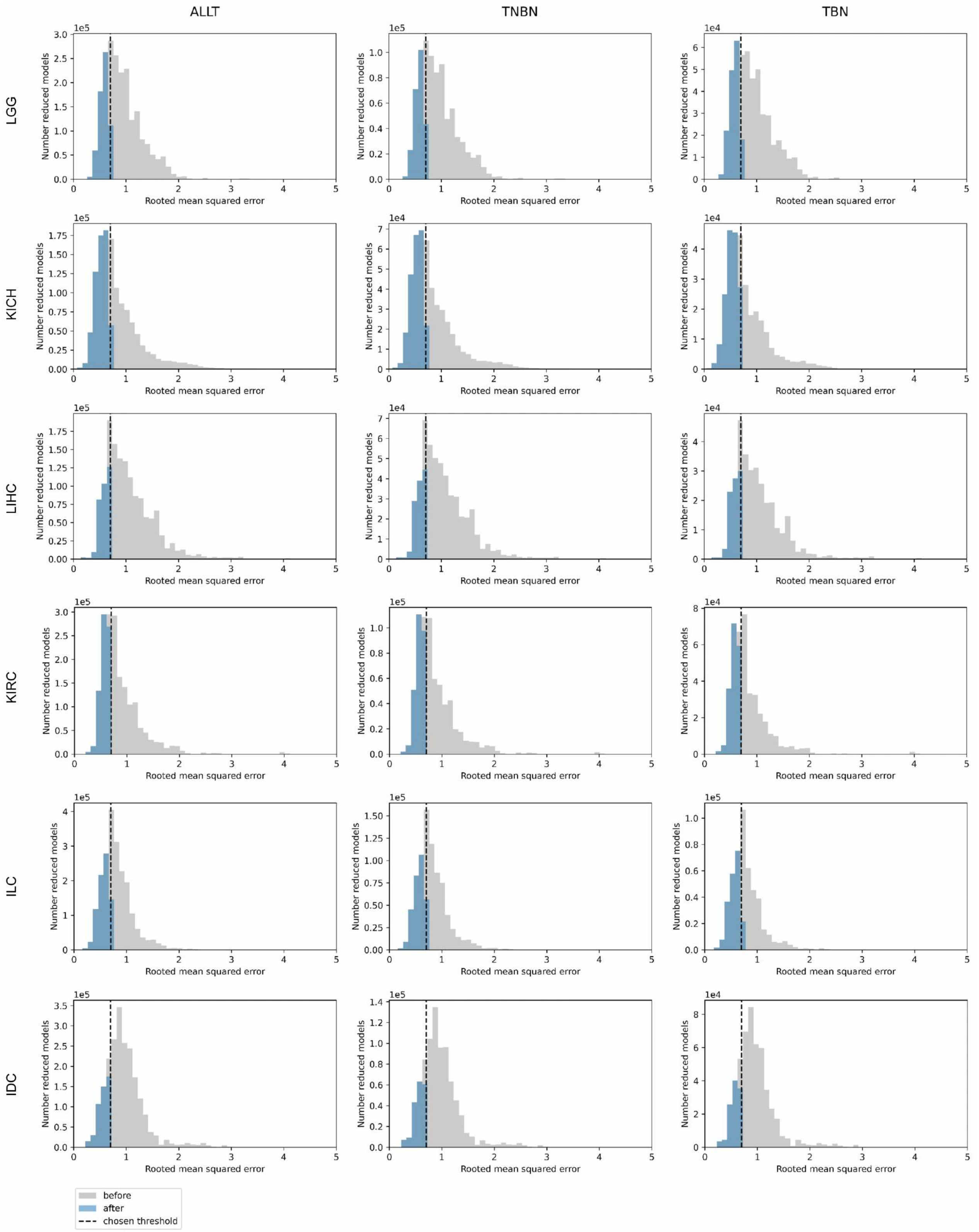


**Figure S2.** Amount of full and reduced models being filtered out by the rooted mean squared error filter depicted for settings ALLT, TNBN, and TBN for Brain lower grade glioma (LGG), Kidney chromophobe carcinoma (KICH), Liver hepatocellular carcinoma (LIHC), Kidney renal cell carcinoma (KIRC) and Breast Invasive Carcinoma types Invasive Lobular Carcinoma (ILC) and Invasive Ductal Carcinoma (IDC).

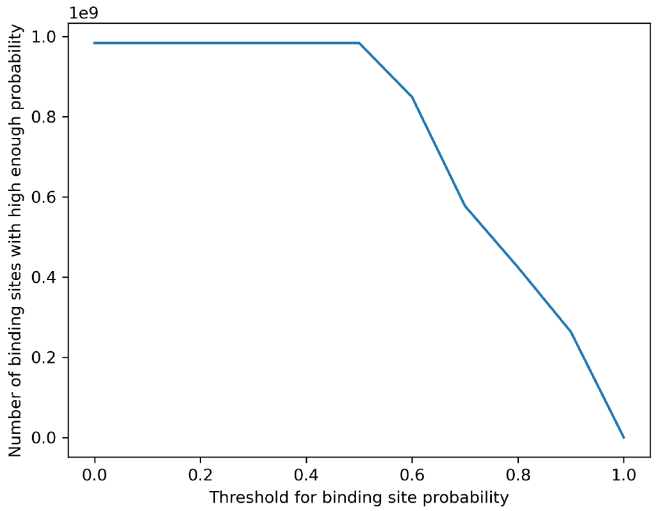


**Figure S3.** Amount of target sites shown for varying binding probability threshold

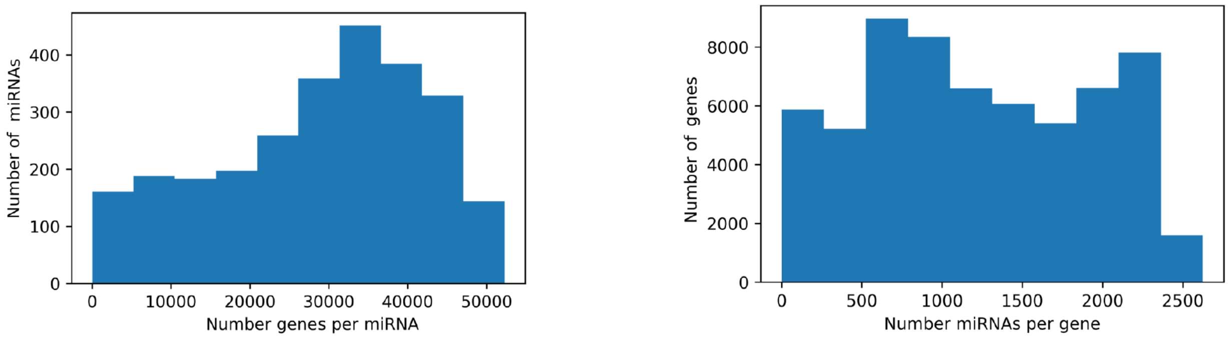


**Figure S4.** Amount of miRNAs per gene and genes per miRNA depicted before any filtering.

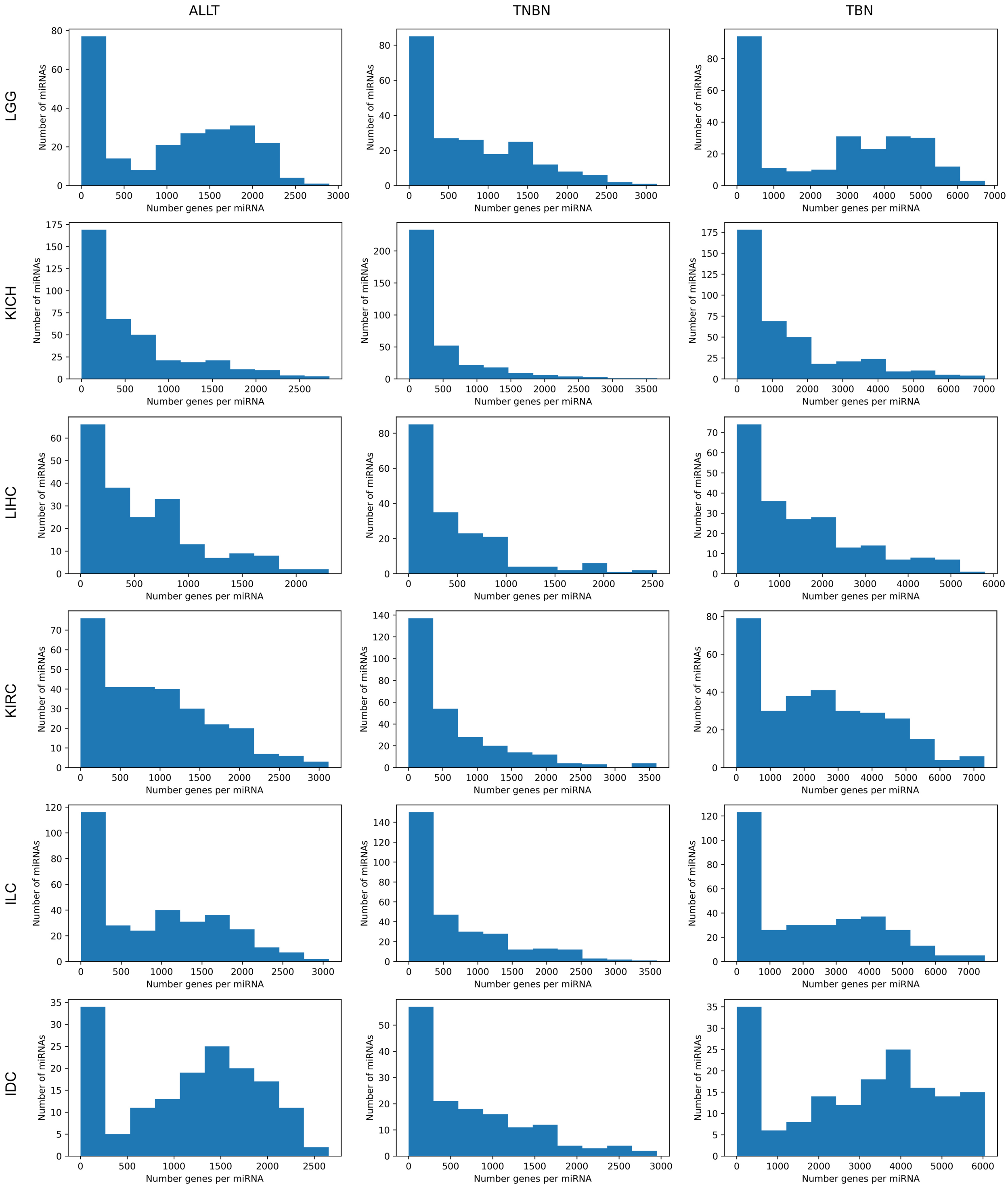
ALLT
TNBN
TBN
LGG
KICH
LIHC
KIRC
ILC
IDC
Number of miRNAs
Number genes per miRNA

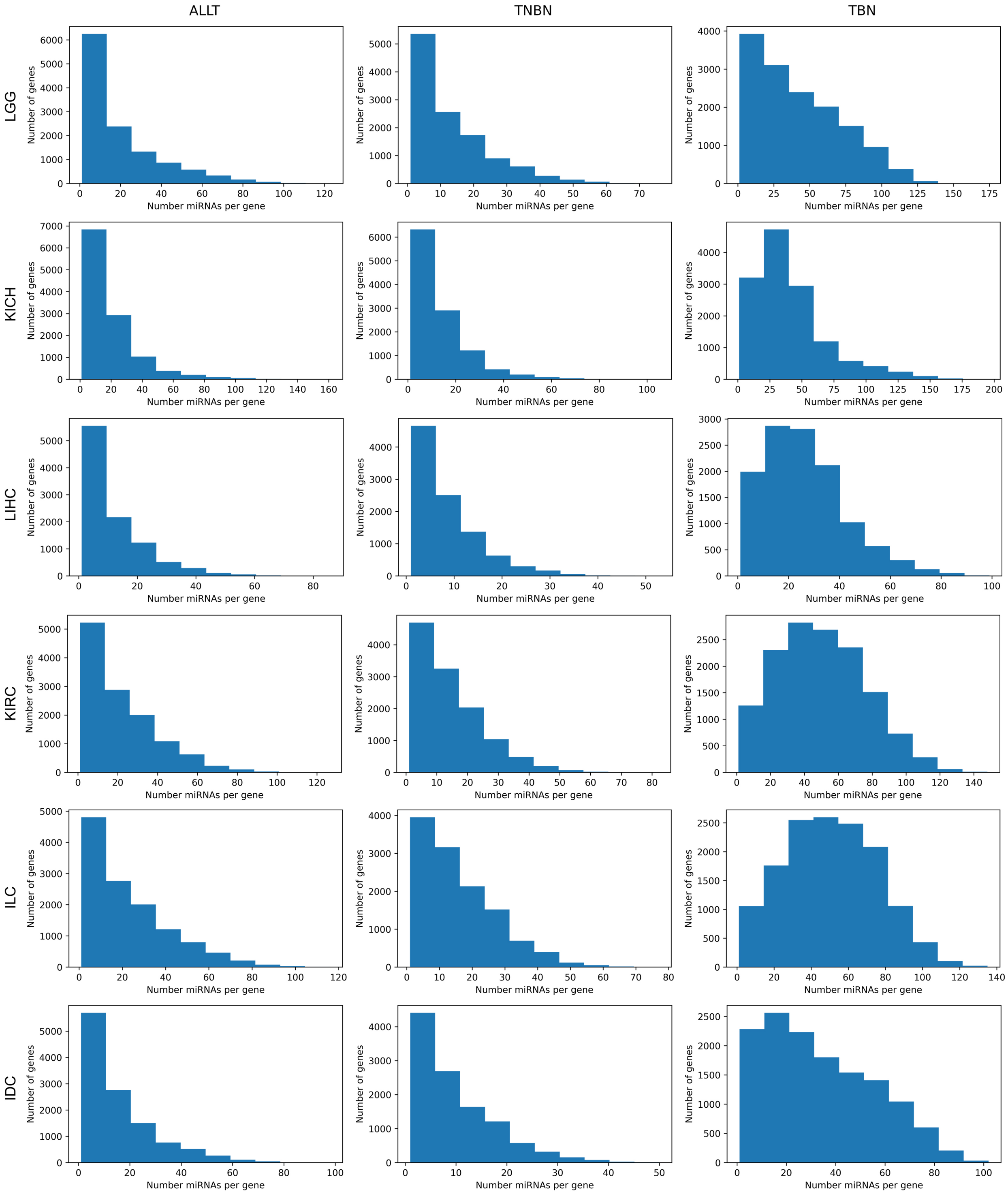


**Figure S5.** Amount of miRNAs per gene and genes per miRNA depicted after filtering for settings ALLT, TNBN, and TBN for Brain lower grade glioma (LGG), Kidney chromophobe carcinoma (KICH), Liver hepatocellular carcinoma (LIHC), Kidney renal cell carcinoma (KIRC) and Breast Invasive Carcinoma types Invasive Lobular Carcinoma (ILC) and Invasive Ductal Carcinoma (IDC).

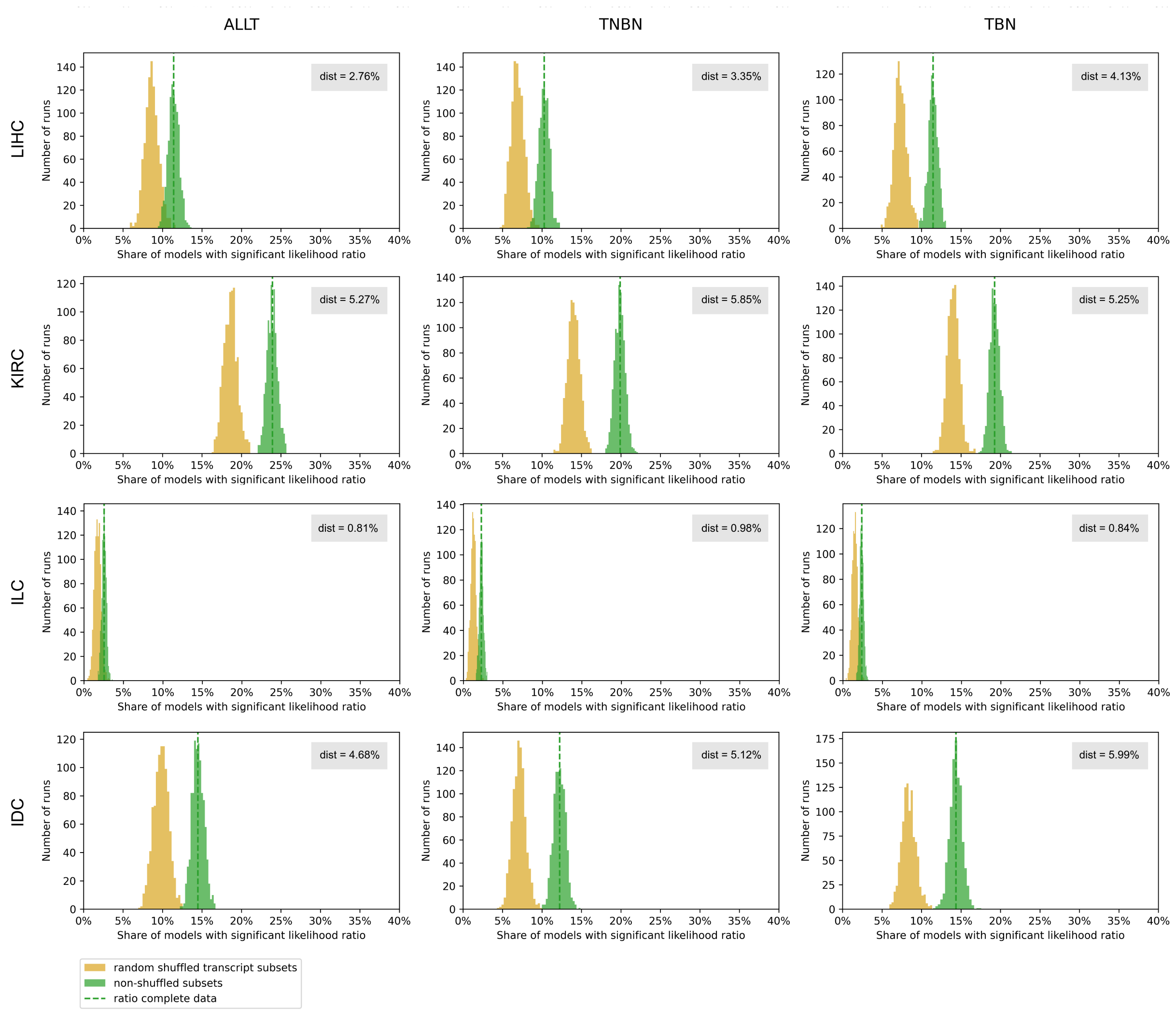


**Figure S6.** The ratio of models with statistically significant (<0.05) corrected p-values of the likelihood ratio test statistic calculated between nested regression models is shown as the dashed green line. To estimate the distribution, the ratio was calculated 1,000 times for random subsets of miRNA-gene pairs (green histogram) and to estimate the impact of alternative splicing, the ratio was calculated 1,000 times for random subsets of miRNA-gene pairs while randomizing the transcript binding labels within a gene (yellow histogram). Dist describes the difference between the average ratio of models based on subsampled real miRNA-gene pairs and after randomizing the transcript category labels. This is shown separately for diseases Liver hepatocellular carcinoma (LIHC), Kidney renal cell carcinoma (KIRC) and Breast Invasive Carcinoma types Invasive Lobular Carcinoma (ILC) and Invasive Ductal Carcinoma (IDC) and for settings ALLT, TNBN, and TBN.

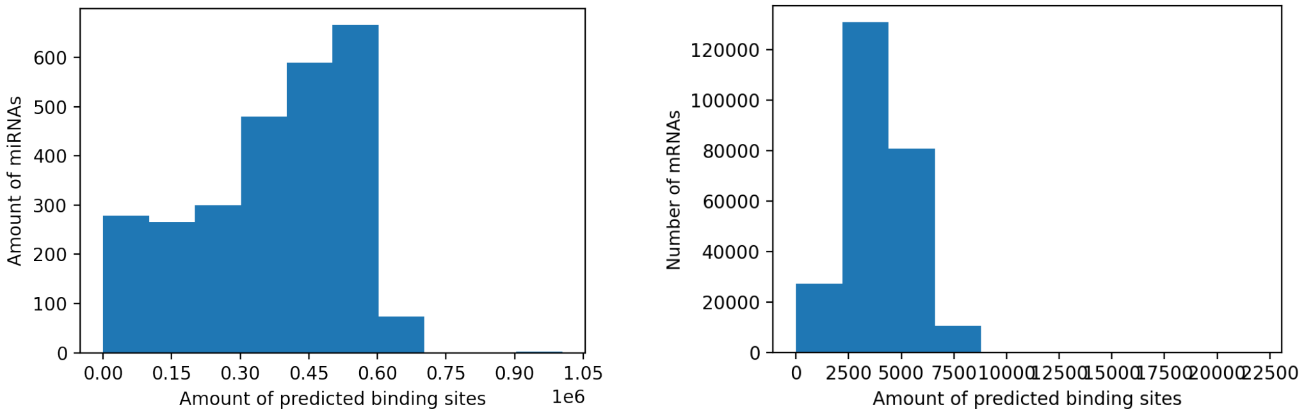


**Figure S7.** Distribution of predicted TarPmiR target sites before any filtering

**Table S2.** Top ten most significantly enriched genesets of non-computational collections (C1, C2, C3, C5, C6, C7, C8, H) for the top 500 genes of miRNA-gene pairs with most significant corrected p-values in LGG setting TNBN.

| Geneset Name | # Genes in Geneset | Description | # Genes in Overlap | FDR q-value |
|---|---|---|---|---|
| TRAVAGLINI_LUNG_PROXIMAL_BASAL_CELL | 629 | - | 63 | 2.06E-34 |
| RUBENSTEIN_SKELETAL_MUSCLE_T_CELLS | 181 | - | 39 | 3E-33 |
| GOMF_RNA_BINDING | 1968 | Binding to an RNA molecule or a portion thereof. [GOC:jl, GOC:mah] | 102 | 3.96E-33 |
| HSIAO_HOUSEKEEPING_GENES | 397 | Housekeeping genes identified as expressed across 19 normal tissues. | 51 | 4.98E-33 |
| BUSSLINGER_DUODENAL_STEM_CELLS | 311 | - | 46 | 2.17E-32 |
| REACTOME_EUKARYOTIC_TRANSLATION_ELONGATION | 94 | Eukaryotic Translation Elongation | 30 | 4.82E-31 |
| REACTOME_METABOLISM_OF_RNA | 714 | Metabolism of RNA | 62 | 4.87E-31 |
| GOCC_SYNAPSE | 1460 | The junction between an axon of one neuron and a dendrite of another neuron, a muscle fiber or a glial cell. As the axon approaches the synapse it enlarges into a specialized structure, the presynaptic terminal bouton, which contains mitochondria and synaptic vesicles. At the tip of the terminal bouton is the presynaptic membrane; facing it, and separated from it by a minute cleft (the synaptic cleft) is a specialized area of membrane on the receiving cell, known as the postsynaptic membrane. In response to the arrival of nerve impulses, the presynaptic terminal bouton secretes molecules of neurotransmitters into the synaptic cleft. These diffuse across the cleft and transmit the signal to the postsynaptic membrane. [GOC:aruk, ISBN:0198506732, PMID:24619342, PMID:29383328, PMID:31998110] | 85 | 7.97E-31 |
| RUBENSTEIN_SKELETAL_MUSCLE_SATELLITE_CELLS | 310 | - | 44 | 2.57E-30 |
| GOBP_CYTOPLASMIC_TRANSLATION | 159 | The chemical reactions and pathways resulting in the formation of a protein in the cytoplasm. This is a ribosome-mediated process in which the information in messenger RNA (mRNA) is used to specify the sequence of amino acids in the protein. [GOC:hjd] | 34 | 3.67E-29 |

**Table S3.** Top ten most significantly enriched genesets of non-computational collections (C1, C2, C3, C5, C6, C7, C8, H) for the top 500 genes of miRNA-gene pairs with most significant corrected p-values in LGG setting TBN.

| Geneset Name | # Genes in Geneset | Description | # Genes in Overlap | FDR q-value |
|---|---|---|---|---|
| BLALOCK_ALZHEIMERS_DISEASE_UP | 1671 | Genes up-regulated in brain from patients with Alzheimer's disease. | 95 | 2.58E-33 |
| BUSSLINGER_GASTRIC_IMMUNE_CELLS | 1492 | - | 76 | 4.68E-23 |
| GRAESSMANN_APOPTOSIS_BY_DOXORUBICIN_DN | 1775 | Genes down-regulated in ME-A cells (breast cancer) undergoing apoptosis in response to doxorubicin [PubChem=31703]. | 83 | 4.68E-23 |
| GOMF_RNA_BINDING | 1968 | Binding to an RNA molecule or a portion thereof. [GOC:jl, GOC:mah] | 84 | 6.63E-21 |
| BUSSLINGER_DUODENAL_IMMUNE_CELLS | 911 | - | 57 | 6.63E-21 |
| SUPT16H_TARGET_GENES | 1979 | Genes containing one or more binding sites for UniProt:Q9Y5B9 (SUPT16H) in their promoter regions (TSS -1000,+100 bp) as identified by GTRD version 20.06 ChIP-seq harmonization. | 83 | 3.03E-20 |
| SCGGAAGY_ELK1_02 | 1242 | Genes having at least one occurrence of the highly conserved motif M3 SCGGAAGY in the regions spanning 4 kb centered on their transcription starting sites [-2kb, +2kb]. This matches the ELK1 [GeneSymbol=ELK1] transcription factor binding site V$ELK1_02 (v7.4 TRANSFAC). | 65 | 3.56E-20 |
| TRAVAGLINI_LUNG_PROXIMAL_BASAL_CELL | 629 | - | 47 | 4.21E-20 |
| GOCC_SYNAPSE | 1460 | The junction between an axon of one neuron and a dendrite of another neuron, a muscle fiber or a glial cell. As the axon approaches the synapse it enlarges into a specialized structure, the presynaptic terminal bouton, which contains mitochondria and synaptic vesicles. At the tip of the terminal bouton is the presynaptic membrane; facing it, and separated from it by a minute cleft (the synaptic cleft) is a specialized area of membrane on the receiving cell, known as the postsynaptic membrane. In response to the arrival of nerve impulses, the presynaptic terminal bouton secretes molecules of neurotransmitters into the synaptic cleft. These diffuse across the cleft and transmit the signal to the postsynaptic membrane. [GOC:aruk, ISBN:0198506732, PMID:24619342, PMID:29383328, PMID:31998110] | 70 | 6.57E-20 |
| MANNO_MIDBRAIN_NEUROTYPES_HOPC | 366 | Cell types are named using anatomical and functional mnemonics prefixed by 'm' or'h' to indicate mouse and human respectively: OMTN, oculomotor and trochlear nucleus; Sert, serotonergic; NbM, medial neuroblast; NbDA, neuroblast dopaminergic; DA0-2, dopaminergic neurons; RN, red nucleus; Gaba1-2, GABAergic neurons; mNbL1-2, lateral neuroblasts; NbML1-5, mediolateral neuroblasts; NProg, neuronal progenitor; Prog, progenitor medial floorplate (FPM), lateral floorplate (FPL), midline (M), basal plate (BP); Rgl1-3, radial glia-like cells; Mgl, microglia; Endo, endothelial cells; Peric, pericytes; Epend, ependymal; OPC, oligodendrocyte precursor cells. | 37 | 7.48E-20 |

**Table S4.** Top ten most significantly enriched genesets of non-computational collections (C1, C2, C3, C5, C6, C7, C8, H) for the top 500 genes of miRNA-gene pairs with most significant corrected p-values in LGG setting ALLT.

| Geneset Name | # Genes in Geneset | Description | # Genes in Overlap | FDR q-value |
|---|---|---|---|---|
| RUBENSTEIN_SKELETAL_MUSCLE_SATELLITE_CELLS | 310 | - | 46 | 7.13E-32 |
| TRAVAGLINI_LUNG_PROXIMAL_BASAL_CELL | 629 | - | 58 | 7.83E-30 |
| GOCC_SYNAPSE | 1460 | The junction between an axon of one neuron and a dendrite of another neuron, a muscle fiber or a glial cell. As the axon approaches the synapse it enlarges into a specialized structure, the presynaptic terminal bouton, which contains mitochondria and synaptic vesicles. At the tip of the terminal bouton is the presynaptic membrane; facing it, and separated from it by a minute cleft (the synaptic cleft) is a specialized area of membrane on the receiving cell, known as the postsynaptic membrane. In response to the arrival of nerve impulses, the presynaptic terminal bouton secretes molecules of neurotransmitters into the synaptic cleft. These diffuse across the cleft and transmit the signal to the postsynaptic membrane. [GOC:aruk, ISBN:0198506732, PMID:24619342, PMID:29383328, PMID:31998110] | 82 | 2.9E-28 |
| GRAESSMANN_APOPTOSIS_BY_DOXORUBICIN_DN | 1775 | Genes down-regulated in ME-A cells (breast cancer) undergoing apoptosis in response to doxorubicin [PubChem=31703]. | 88 | 8.49E-27 |
| BUSSLINGER_DUODENAL_STEM_CELLS | 311 | - | 41 | 1.31E-26 |
| RUBENSTEIN_SKELETAL_MUSCLE_T_CELLS | 181 | - | 33 | 1.1E-25 |
| REACTOME_INFECTIOUS_DISEASE | 1019 | Infectious disease | 62 | 1.24E-22 |
| BUSSLINGER_DUODENAL_DIFFERENTIATING_STEM_CELLS | 305 | - | 37 | 1.24E-22 |
| SETD1A_TARGET_GENES | 1531 | Genes containing one or more binding sites for UniProt:O15047 (SETD1A) in their promoter regions (TSS -1000,+100 bp) as identified by GTRD version 20.06 ChIP-seq harmonization. | 75 | 2.97E-22 |
| BLALOCK_ALZHEIMERS_DISEASE_UP | 1671 | Genes up-regulated in brain from patients with Alzheimer's disease. | 78 | 5.03E-22 |

**Table S5.** Top ten most significantly enriched genesets of non-computational collections (C1, C2, C3, C5, C6, C7, C8, H) for the top 500 genes of miRNA-gene pairs with most significant corrected p-values in LIHC setting TNBN.

| Geneset Name | # Genes in Geneset | Description | # Genes in Overlap | FDR q-value |
|---|---|---|---|---|
| HSIAO_LIVER_SPECIFIC_GENES | 249 | Liver selective genes | 50 | 1.21E-41 |
| GOMF_RNA_BINDING | 1968 | Binding to an RNA molecule or a portion thereof. [GOC:jl, GOC:mah] | 108 | 1.63E-37 |
| PUJANA_BRCA1_PCC_NETWORK | 1625 | Genes constituting the BRCA1-PCC network of transcripts whose expression positively correlated (Pearson correlation coefficient, PCC >= 0.4) with that of BRCA1 [GeneID=672] across a compendium of normal tissues. | 97 | 2.62E-36 |
| AIZARANI_LIVER_C11_HEPATOCYTES_1 | 298 | - | 48 | 1.23E-35 |
| AIZARANI_LIVER_C14_HEPATOCYTES_2 | 226 | - | 43 | 6.33E-35 |
| REACTOME_METABOLISM_OF_RNA | 714 | Metabolism of RNA | 63 | 6.22E-32 |
| CAIRO_HEPATOBLASTOMA_CLASSES_UP | 612 | Genes up-regulated in robust Cluster 2 (rC2) of hepatoblastoma samples compared to those in the robust Cluster 1 (rC1). | 59 | 7.01E-32 |
| GRAESSMANN_APOPTOSIS_BY_DOXORUBICIN_DN | 1775 | Genes down-regulated in ME-A cells (breast cancer) undergoing apoptosis in response to doxorubicin [PubChem=31703]. | 90 | 2.47E-28 |
| GOCC_NUCLEAR_PROTEIN_CONTAINING_COMPLEX | 1243 | A stable assembly of two or more macromolecules, i.e. proteins, nucleic acids, carbohydrates or lipids, in which at least one component is a protein and the constituent parts function together in the nucleus. [GOC:pg] | 73 | 3.27E-26 |
| TRAVAGLINI_LUNG_PROLIFERATING_BASAL_CELL | 891 | - | 62 | 8.72E-26 |

**Table S6.** Top ten most significantly enriched genesets of non-computational collections (C1, C2, C3, C5, C6, C7, C8, H) for the top 500 genes of miRNA-gene pairs with most significant corrected p-values in LIHC setting TBN.

| Geneset Name | # Genes in Geneset | Description | # Genes in Overlap | FDR q-value |
|---|---|---|---|---|
| CARRILLOREIXACH_HEPATOBLASTOMA_VS_NORMAL_DN | 1259 | Genes down-regulated in hepatoblastoma (HB) tumors as compared with non-tumor (NT) adjacent tissue. | 105 | 4E-53 |
| GOBP_SMALL_MOLECULE_METABOLIC_PROCESS | 1830 | The chemical reactions and pathways involving small molecules, any low molecular weight, monomeric, non-encoded molecule. [GOC:curators, GOC:pde, GOC:vw] | 117 | 4.6E-48 |
| HOSHIDA_LIVER_CANCER_SUBCLASS_S3 | 266 | Genes from 'subtype S3' signature of hepatocellular carcinoma (HCC): hepatocyte differentiation. | 52 | 2.4E-43 |
| HSIAO_LIVER_SPECIFIC_GENES | 249 | Liver selective genes | 49 | 6.38E-41 |
| AIZARANI_LIVER_C11_HEPATOCYTES_1 | 298 | - | 51 | 1.38E-39 |
| GOBP_ORGANIC_ACID_METABOLIC_PROCESS | 969 | The chemical reactions and pathways involving organic acids, any acidic compound containing carbon in covalent linkage. [ISBN:0198506732] | 77 | 1.06E-36 |
| AIZARANI_LIVER_C14_HEPATOCYTES_2 | 226 | - | 42 | 8.88E-34 |
| CHIANG_LIVER_CANCER_SUBCLASS_PROLIFERATION_DN | 179 | Top 200 marker genes down-regulated in the 'proliferation' subclass of hepatocellular carcinoma (HCC); characterized by increased proliferation, high levels of serum AFP [GeneID=174], and chromosomal instability. | 36 | 6.23E-30 |
| GOCC_MITOCHONDRION | 1672 | A semiautonomous, self replicating organelle that occurs in varying numbers, shapes, and sizes in the cytoplasm of virtually all eukaryotic cells. It is notably the site of tissue respiration. [GOC:giardia, ISBN:0198506732] | 87 | 3.55E-28 |
| FLECHNER_BIOPSY_KIDNEY_TRANSPLANT_REJECTED_VS_OK_DN | 553 | Genes down-regulated in kidney biopsies from patients with acute transplant rejection compared to the biopsies from patients with well functioning kidneys more than 1-year post transplant. | 52 | 2.12E-27 |

**Table S7.** Top ten most significantly enriched genesets of non-computational collections (C1, C2, C3, C5, C6, C7, C8, H) for the top 500 genes of miRNA-gene pairs with most significant corrected p-values in LIHC setting ALLT.

| Geneset Name | # Genes in Geneset | Description | # Genes in Overlap | FDR q-value |
|---|---|---|---|---|
| GOMF_RNA_BINDING | 1968 | Binding to an RNA molecule or a portion thereof. [GOC:jl, GOC:mah] | 115 | 2.98E-43 |
| PUJANA_BRCA1_PCC_NETWORK | 1625 | Genes constituting the BRCA1-PCC network of transcripts whose expression positively correlated (Pearson correlation coefficient, PCC $>=$ 0.4) with that of BRCA1 [GeneID=672] across a compendium of normal tissues. | 103 | 1.68E-41 |
| CAIRO_HEPATOBLASTOMA_CLASSES_UP | 612 | Genes up-regulated in robust Cluster 2 (rC2) of hepatoblastoma samples compared to those in the robust Cluster 1 (rC1). | 66 | 4.82E-39 |
| REACTOME_METABOLISM_OF_RNA | 714 | Metabolism of RNA | 66 | 5.76E-35 |
| DODD_NASOPHARYNGEAL_CARCINOMA_DN | 1405 | Genes down-regulated in nasopharyngeal carcinoma (NPC) compared to the normal tissue. | 87 | 8.25E-34 |
| ELF2_TARGET_GENES | 1511 | Genes containing one or more binding sites for UniProt:Q15723 (ELF2) in their promoter regions (TSS -1000,+100 bp) as identified by GTRD version 20.06 ChIP-seq harmonization. | 87 | 1.65E-31 |
| HE_LIM_SUN_FETAL_LUNG_C5_LARGE_PRE_B_CELL | 1363 | Large pre-B | 80 | 2.87E-29 |
| HOUNKPE_HOUSEKEEPING_GENES | 1129 | List of 1130 human and mouse housekeeping genes. This list shows the overlap of human genes stably expressed across 52 tissues and cells types and mouse genes with at least one of their transcripts expressed across 14 tissues and cells types. | 71 | 1.91E-27 |
| GOCC_NUCLEAR_PROTEIN_CONTAINING_COMPLEX | 1243 | A stable assembly of two or more macromolecules, i.e. proteins, nucleic acids, carbohydrates or lipids, in which at least one component is a protein and the constituent parts function together in the nucleus. [GOC:pg] | 73 | 1.57E-26 |
| FISCHER_DREAM_TARGETS | 969 | Target genes of the DREAM complex. | 65 | 1.57E-26 |

**Table S8.** Top ten most significantly enriched genesets of non-computational collections (C1, C2, C3, C5, C6, C7, C8, H) for the top 500 genes of miRNA-gene pairs with most significant corrected p-values in KIRC setting TNBN.

| Geneset Name | # Genes in Geneset | Description | # Genes in Overlap | FDR q-value |
|---|---|---|---|---|
| MURARO_PANCREAS_DUCTAL_CELL | 1276 | - | 82 | 8.39E-32 |
| BUSSLINGER_GASTRIC_IMMUNE_CELLS | 1492 | - | 75 | 3.54E-22 |
| HP_ABNORMAL_CIRCULATING_METABOLITE_CONCENTRATION | 1283 | Abnormal circulating metabolite concentration | 68 | 3.23E-21 |
| GOBP_CELL_ADHESION | 1524 | The attachment of a cell, either to another cell or to an underlying substrate such as the extracellular matrix, via cell adhesion molecules. [GOC:hb, GOC:pf] | 74 | 3.23E-21 |
| GOBP_CYTOSKELETON_ORGANIZATION | 1509 | A process that is carried out at the cellular level which results in the assembly, arrangement of constituent parts, or disassembly of cytoskeletal structures. [GOC:dph, GOC:jl, GOC:mah] | 71 | 1.62E-19 |
| BLALOCK_ALZHEIMERS_DISEASE_UP | 1671 | Genes up-regulated in brain from patients with Alzheimer's disease. | 74 | 5.1E-19 |
| GRAESSMANN_APOPTOSIS_BY_DOXORUBICIN_DN | 1775 | Genes down-regulated in ME-A cells (breast cancer) undergoing apoptosis in response to doxorubicin [PubChem=31703]. | 75 | 3.43E-18 |
| LAKE_ADULT_KIDNEY_C3_PROXIMAL_TUBULE_EPITHELIAL_CELLS_S1_S2 | 221 | - | 29 | 3.8E-18 |
| GOBP_SMALL_MOLECULE_METABOLIC_PROCESS | 1830 | The chemical reactions and pathways involving small molecules, any low molecular weight, monomeric, non-encoded molecule. [GOC:curators, GOC:pde, GOC:vw] | 76 | 3.8E-18 |
| NAKAYA_PBMC_FLUMIST_AGE_18_50YO_7DY_UP | 1575 | Genes up-regulated in peripheral blood mononuclear cell 7d vs 0d in adults (18-50) after exposure to FluMist , time point 7D. Comment: Supplementary Table 1b: All the differentially expressed genes identified in PBMCs of TIV vaccinees. | 70 | 4.12E-18 |

**Table S9.** Top ten most significantly enriched genesets of non-computational collections (C1, C2, C3, C5, C6, C7, C8, H) for the top 500 genes of miRNA-gene pairs with most significant corrected p-values in KIRC setting TBN.

| Geneset Name | # Genes in Geneset | Description | # Genes in Overlap | FDR q-value |
|---|---|---|---|---|
| MURARO_PANCREAS_DUCTAL_CELL | 1276 | - | 77 | 9.81E-28 |
| BUSSLINGER_GASTRIC_IMMUNE_CELLS | 1492 | - | 79 | 4.56E-25 |
| GOBERT_OLIGODENDROCYTE_DIFFERENTIATION_DN | 1088 | Genes down-regulated during differentiation of Oli-Neu cells (oligodendroglial precursor) in response to PD174265 [PubChem=4709]. | 64 | 3.52E-22 |
| NAKAYA_PBMC_FLUMIST_AGE_18_50YO_7DY_UP | 1575 | Genes up-regulated in peripheral blood mononuclear cell 7d vs 0d in adults (18-50) after exposure to FluMist , time point 7D. Comment: Supplementary Table 1b: All the differentially expressed genes identified in PBMCs of TIV vaccinees. | 71 | 2.3E-18 |
| DODD_NASOPHARYNGEAL_CARCINOMA_UP | 1813 | Genes up-regulated in nasopharyngeal carcinoma (NPC) compared to the normal tissue. | 76 | 3.43E-18 |
| GOCC_ANCHORING_JUNCTION | 899 | A cell junction that mechanically attaches a cell (and its cytoskeleton) to neighboring cells or to the extracellular matrix. [ISBN:0815332181] | 53 | 3.43E-18 |
| GRAESSMANN_APOPTOSIS_BY_DOXORUBICIN_DN | 1775 | Genes down-regulated in ME-A cells (breast cancer) undergoing apoptosis in response to doxorubicin [PubChem=31703]. | 75 | 3.43E-18 |
| MIR8485 | 1034 | Genes predicted to be targets of miRBase v22 microRNA hsa-miR-8485 in miRDB v6.0 with MirTarget v4 prediction scores > 80 (high confidence targets). | 56 | 1.05E-17 |
| GOBP_SMALL_MOLECULE_METABOLIC_PROCESS | 1830 | The chemical reactions and pathways involving small molecules, any low molecular weight, monomeric, non-encoded molecule. [GOC:curators, GOC:pde, GOC:vw] | 74 | 6.38E-17 |
| MANNO_MIDBRAIN_NEUROTYPES_HENDO | 888 | Cell types are named using anatomical and functional mnemonics prefixed by 'm' or'h' to indicate mouse and human respectively: OMTN, oculomotor and trochlear nucleus; Sert, serotonergic; NbM, medial neuroblast; NbDA, neuroblast dopaminergic; DA0-2, dopaminergic neurons; RN, red nucleus; Gaba1-2, GABAergic neurons; mNbL1-2, lateral neuroblasts; NbML1-5, mediolateral neuroblasts; NProg, neuronal progenitor; Prog, progenitor medial floorplate (FPM), lateral floorplate (FPL), midline (M), basal plate (BP); Rgl1-3, radial glia-like cells; Mgl, microglia; Endo, endothelial cells; Peric, pericytes; Epend, ependymal; OPC, oligodendrocyte precursor cells. | 50 | 2.59E-16 |

**Table S10.** Top ten most significantly enriched genesets of non-computational collections (C1, C2, C3, C5, C6, C7, C8, H) for the top 500 genes of miRNA-gene pairs with most significant corrected p-values in KIRC setting ALLT.

| Geneset Name | # Genes in Geneset | Description | # Genes in Overlap | FDR q-value |
|---|---|---|---|---|
| MURARO_PANCREAS_DUCTAL_CELL | 1276 | - | 80 | 3.22E-30 |
| KRIEG_HYPOXIA_NOT_VIA_KDM3A | 746 | Genes induced under hypoxia independently of KDM3A [GeneID=55818] in RCC4 cells (renal carcinoma) expressing VHL [GeneID=7428]. | 55 | 3.7E-23 |
| BUSSLINGER_GASTRIC_IMMUNE_CELLS | 1492 | - | 70 | 5.96E-19 |
| GOBP_REGULATION_OF_INTRACELLULAR_SIGNAL_TRANSDUCTION | 1770 | Any process that modulates the frequency, rate or extent of intracellular signal transduction. [GOC:dph, GOC:signaling, GOC:tb, GOC:TermGenie] | 74 | 1.49E-17 |
| TRAVAGLINI_LUNG_PROXIMAL_CILIATED_CELL | 1770 | - | 74 | 1.49E-17 |
| GRAESSMANN_APOPTOSIS_BY_DOXORUBICIN_DN | 1775 | Genes down-regulated in ME-A cells (breast cancer) undergoing apoptosis in response to doxorubicin [PubChem=31703]. | 74 | 1.49E-17 |
| ONKEN_UVEAL_MELANOMA_UP | 790 | Genes up-regulated in uveal melanoma: class 2 vs class 1 tumors. | 48 | 8.76E-17 |
| GOBP_SMALL_MOLECULE_METABOLIC_PROCESS | 1830 | The chemical reactions and pathways involving small molecules, any low molecular weight, monomeric, non-encoded molecule. [GOC:curators, GOC:pde, GOC:vw] | 73 | 2.55E-16 |
| ZNF768_TARGET_GENES | 1362 | Genes containing one or more binding sites for UniProt:Q9H5H4 (ZNF768) in their promoter regions (TSS -1000,+100 bp) as identified by GTRD version 20.06 ChIP-seq harmonization. | 62 | 3.33E-16 |
| GRYDER_PAX3FOXO1_ENHANCERS_IN_TADS | 1009 | Expressed genes (FPKM>1) associated with high-confidence PAX3-FOXO1 sites with enhancers in primary tumors and cell lines, restricted to those within topological domain boundaries | 53 | 3.71E-16 |

**Table S11.** Top ten most significantly enriched genesets of non-computational collections (C1, C2, C3, C5, C6, C7, C8, H) for the top 500 genes of miRNA-gene pairs with most significant corrected p-values in KICH setting TNBN.

| Geneset Name | # Genes in Geneset | Description | # Genes in Overlap | FDR q-value |
|---|---|---|---|---|
| CHARAFE_BREAST_CANCER_LUMINAL_VS_MESENCHYMAL_UP | 454 | Genes up-regulated in luminal-like breast cancer cell lines compared to the mesenchymal-like ones. | 52 | 1.08E-30 |
| TRAVAGLINI_LUNG_PROXIMAL_CILIATED_CELL | 1770 | - | 87 | 8.31E-26 |
| GOCC_MITOCHONDRION | 1672 | A semiautonomous, self replicating organelle that occurs in varying numbers, shapes, and sizes in the cytoplasm of virtually all eukaryotic cells. It is notably the site of tissue respiration. [GOC:giardia, ISBN:0198506732] | 82 | 3.62E-24 |
| DODD_NASOPHARYNGEAL_CARCINOMA_UP | 1813 | Genes up-regulated in nasopharyngeal carcinoma (NPC) compared to the normal tissue. | 84 | 2.66E-23 |
| MURARO_PANCREAS_DUCTAL_CELL | 1276 | - | 69 | 2.09E-22 |
| BLALOCK_ALZHEIMERS_DISEASE_UP | 1671 | Genes up-regulated in brain from patients with Alzheimer's disease. | 79 | 2.09E-22 |
| CHARAFE_BREAST_CANCER_LUMINAL_VS_BASAL_UP | 384 | Genes up-regulated in luminal-like breast cancer cell lines compared to the basal-like ones. | 40 | 4.96E-22 |
| LIM_MAMMARY_STEM_CELL_DN | 416 | Genes consistently down-regulated in mammary stem cells both in mouse and human species. | 41 | 9.02E-22 |
| GOBP_CYTOSKELETON_ORGANIZATION | 1509 | A process that is carried out at the cellular level which results in the assembly, arrangement of constituent parts, or disassembly of cytoskeletal structures. [GOC:dph, GOC:jl, GOC:mah] | 71 | 8E-20 |
| STEIN_ESRRA_TARGETS_UP | 385 | Genes up-regulated by ESRRA [GeneID=2101] only. | 37 | 4.08E-19 |

**Table S12.** Top ten most significantly enriched genesets of non-computational collections (C1, C2, C3, C5, C6, C7, C8, H) for the top 500 genes of miRNA-gene pairs with most significant corrected p-values in KICH setting TBN.

| Geneset Name | # Genes in Geneset | Description | # Genes in Overlap | FDR q-value |
|---|---|---|---|---|
| RODRIGUES_THYROID_CARCINOMA_POORLY_DIFFERENTIATED_DN | 802 | Genes down-regulated in poorly differentiated thyroid carcinoma (PDTC) compared to normal thyroid tissue. | 61 | 1.98E-26 |
| LAKE_ADULT_KIDNEY_C20_COLLECTING_DUCT_INTERCALATED_CELLS_TYPE_A_CORTEX | 152 | - | 32 | 2.24E-26 |
| CHARAFE_BREAST_CANCER_LUMINAL_VS_MESENCHYMAL_UP | 454 | Genes up-regulated in luminal-like breast cancer cell lines compared to the mesenchymal-like ones. | 44 | 9.35E-23 |
| GOBP_SMALL_MOLECULE_METABOLIC_PROCESS | 1830 | The chemical reactions and pathways involving small molecules, any low molecular weight, monomeric, non-encoded molecule. [GOC:curators, GOC:pde, GOC:vw] | 81 | 6.42E-21 |
| STEIN_ESRRA_TARGETS | 529 | Genes regulated by ESRRA [GeneID=2101] in MCF-7 cells (breast cancer). | 44 | 2.89E-20 |
| LIM_MAMMARY_STEM_CELL_DN | 416 | Genes consistently down-regulated in mammary stem cells both in mouse and human species. | 39 | 1.33E-19 |
| STEIN_ESRRA_TARGETS_UP | 385 | Genes up-regulated by ESRRA [GeneID=2101] only. | 37 | 6.69E-19 |
| MURARO_PANCREAS_DUCTAL_CELL | 1276 | - | 64 | 7.48E-19 |
| TGACCTY_ERR1_Q2 | 1064 | Genes having at least one occurrence of the highly conserved motif M25 TGACCTY in the regions spanning 4 kb centered on their transcription starting sites [-2kb, +2kb]. This matches the ESRRA [GeneSymbol=ESRRA] transcription factor binding site V$ERR1_Q2 (v7.4 TRANSFAC). | 58 | 1.38E-18 |
| LAKE_ADULT_KIDNEY_C21_COLLECTING_DUCT_INTERCALATED_CELLS_TYPE_B | 103 | - | 22 | 4.65E-18 |

**Table S13.** Top ten most significantly enriched genesets of non-computational collections (C1, C2, C3, C5, C6, C7, C8, H) for the top 500 genes of miRNA-gene pairs with most significant corrected p-values in KICH setting ALLT.

| Geneset Name | # Genes in Geneset | Description | # Genes in Overlap | FDR q-value |
|---|---|---|---|---|
| CHARAFE_BREAST_CANCER_LUMINAL_VS_MESENCHYMAL_UP | 454 | Genes up-regulated in luminal-like breast cancer cell lines compared to the mesenchymal-like ones. | 47 | 2.14E-25 |
| RODRIGUES_THYROID_CARCINOMA_POORLY_DIFFERENTIATED_DN | 802 | Genes down-regulated in poorly differentiated thyroid carcinoma (PDTC) compared to normal thyroid tissue. | 58 | 3.49E-24 |
| LAKE_ADULT_KIDNEY_C12_THICK_ASCENDING_LIMB | 381 | - | 41 | 7.82E-23 |
| CHEMNITZ_RESPONSE_TO_PROSTAGLANDIN_E2_DN | 408 | Genes down-regulated in CD4+ [GeneID=920] T lymphocytes after stimulation with prostaglandin E2 [PubChem=5280360]. | 39 | 8.52E-20 |
| LIM_MAMMARY_STEM_CELL_DN | 416 | Genes consistently down-regulated in mammary stem cells both in mouse and human species. | 39 | 1.39E-19 |
| MURARO_PANCREAS_DUCTAL_CELL | 1276 | - | 64 | 8.03E-19 |
| TRAVAGLINI_LUNG_PROXIMAL_CILIATED_CELL | 1770 | - | 75 | 2.57E-18 |
| DODD_NASOPHARYNGEAL_CARCINOMA_UP | 1813 | Genes up-regulated in nasopharyngeal carcinoma (NPC) compared to the normal tissue. | 74 | 3.75E-17 |
| MENON_FETAL_KIDNEY_8_CONNECTING_TUBULE_CELLS | 267 | - | 30 | 5.96E-17 |
| COLDREN_GEFITINIB_RESISTANCE_DN | 229 | Genes down-regulated in NSCLC (non-small cell lung carcinoma) cell lines resistant to gefitinib [PubChem=123631] compared to the sensitive ones. | 28 | 1.04E-16 |

**Table S14.** Top ten most significantly enriched genesets of non-computational collections (C1, C2, C3, C5, C6, C7, C8, H) for the top 500 genes of miRNA-gene pairs with most significant corrected p-values in ILC setting TNBN.

| Geneset Name | # Genes in Geneset | Description | # Genes in Overlap | FDR q-value |
|---|---|---|---|---|
| DIAZ_CHRONIC_MYELOGENOUS_LEUKEMIA_UP | 1399 | Genes up-regulated in CD34+ [GeneID=947] cells isolated from bone marrow of CML (chronic myelogenous leukemia) patients, compared to those from normal donors. | 79 | 1.45E-26 |
| PUJANA_BRCA1_PCC_NETWORK | 1625 | Genes constituting the BRCA1-PCC network of transcripts whose expression positively correlated (Pearson correlation coefficient, PCC >= 0.4) with that of BRCA1 [GeneID=672] across a compendium of normal tissues. | 79 | 1.37E-22 |
| BLALOCK_ALZHEIMERS_DISEASE_UP | 1671 | Genes up-regulated in brain from patients with Alzheimer's disease. | 79 | 5.48E-22 |
| HP_PEDIATRIC_ONSET | 1592 | Pediatric onset | 76 | 2.21E-21 |
| HP_ABNORMALITY_OF_THE_FOREHEAD | 1128 | Abnormality of the forehead | 63 | 9.53E-21 |
| HP_ABNORMAL_JOINT_MORPHOLOGY | 1151 | Abnormal joint morphology | 63 | 2.31E-20 |
| HP_ABNORMALITY_OF_THE_CURVATURE_OF_THE_VERTEBRAL_COLUMN | 1136 | Abnormality of the curvature of the vertebral column | 62 | 5.45E-20 |
| HP_ABNORMALITY_OF_THE_OUTER_EAR | 1230 | Abnormality of the outer ear | 64 | 1.03E-19 |
| RODRIGUES_THYROID_CARCINOMA_POORLY_DIFFERENTIATED_DN | 802 | Genes down-regulated in poorly differentiated thyroid carcinoma (PDTC) compared to normal thyroid tissue. | 52 | 1.03E-19 |
| HP_ABNORMAL_NASAL_BRIDGE_MORPHOLOGY | 1039 | Abnormal nasal bridge morphology | 58 | 3.92E-19 |

**Table S15.** Top ten most significantly enriched genesets of non-computational collections (C1, C2, C3, C5, C6, C7, C8, H) for the top 500 genes of miRNA-gene pairs with most significant corrected p-values in ILC setting TBN.

| Geneset Name | # Genes in Geneset | Description | # Genes in Overlap | FDR q-value |
|---|---|---|---|---|
| GRAESSMANN_APOPTOSIS_BY_DOXORUBICIN_DN | 1775 | Genes down-regulated in ME-A cells (breast cancer) undergoing apoptosis in response to doxorubicin [PubChem=31703]. | 89 | 7.61E-27 |
| NAKAYA_PBMC_FLUMIST_AGE_18_50YO_7DY_UP | 1575 | Genes up-regulated in peripheral blood mononuclear cell 7d vs 0d in adults (18-50) after exposure to FluMist , time point 7D. Comment: Supplementary Table 1b: All the differentially expressed genes identified in PBMCs of TIV vaccinees. | 75 | 8.56E-21 |
| DIAZ_CHRONIC_MYELOGENOUS_LEUKEMIA_UP | 1399 | Genes up-regulated in CD34+ [GeneID=947] cells isolated from bone marrow of CML (chronic myelogenous leukemia) patients, compared to those from normal donors. | 67 | 1.83E-18 |
| GOMF_RNA_BINDING | 1968 | Binding to an RNA molecule or a portion thereof. [GOC:jl, GOC:mah] | 80 | 2.05E-18 |
| BLALOCK_ALZHEIMERS_DISEASE_UP | 1671 | Genes up-regulated in brain from patients with Alzheimer's disease. | 73 | 2.4E-18 |
| GOBP_CELL_PROJECTION_ORGANIZATION | 1576 | A process that is carried out at the cellular level which results in the assembly, arrangement of constituent parts, or disassembly of a prolongation or process extending from a cell, e.g. a flagellum or axon. [GOC:jl, GOC:mah, http://www.cogsci.princeton.edu/ wn/] | 70 | 6.34E-18 |
| GOBP_POSITIVE_REGULATION_OF_CELLULAR_COMPONENT_ORGANIZATION | 1068 | Any process that activates or increases the frequency, rate or extent of a process involved in the formation, arrangement of constituent parts, or disassembly of cell structures, including the plasma membrane and any external encapsulating structures such as the cell wall and cell envelope. [GOC:ai] | 57 | 9.35E-18 |
| SALL4_TARGET_GENES | 1870 | Genes containing one or more binding sites for UniProt:Q9UJQ4 (SALL4) in their promoter regions (TSS -1000,+100 bp) as identified by GTRD version 20.06 ChIP-seq harmonization. | 76 | 1.34E-17 |
| HP_ABNORMAL_CARDIAC_SEPTUM_MORPHOLOGY | 734 | Abnormal cardiac septum morphology | 47 | 2.07E-17 |
| NAKAYA_PLASMACYTOID_DENDRITIC_CELL_FLUMIST_AGE_18_50YO_7DY_UP | 1215 | Genes up-regulated in plasmacytoid dendritic cell 7d vs 0d in young adults (18-50) after exposure to FluMist , time point 7D | 60 | 2.44E-17 |

**Table S16.** Top ten most significantly enriched genesets of non-computational collections (C1, C2, C3, C5, C6, C7, C8, H) for the top 500 genes of miRNA-gene pairs with most significant corrected p-values in ILC setting ALLT.

| Geneset Name | # Genes in Geneset | Description | # Genes in Overlap | FDR q-value |
|---|---|---|---|---|
| DIAZ_CHRONIC_MYELOGENOUS_LEUKEMIA_UP | 1399 | Genes up-regulated in CD34+ [GeneID=947] cells isolated from bone marrow of CML (chronic myelogenous leukemia) patients, compared to those from normal donors. | 69 | 1.62E-19 |
| PUJANA_BRCA1_PCC_NETWORK | 1625 | Genes constituting the BRCA1-PCC network of transcripts whose expression positively correlated (Pearson correlation coefficient, PCC >= 0.4) with that of BRCA1 [GeneID=672] across a compendium of normal tissues. | 73 | 7.9E-19 |
| HE_LIM_SUN_FETAL_LUNG_C2_HSC_CELL | 1635 | HSC | 73 | 7.9E-19 |
| GOMF_RNA_BINDING | 1968 | Binding to an RNA molecule or a portion thereof. [GOC:jl, GOC:mah] | 80 | 1.39E-18 |
| GOBP_INTRACELLULAR_TRANSPORT | 1591 | The directed movement of substances within a cell. [GOC:ai] | 71 | 2.03E-18 |
| GRAESSMANN_APOPTOSIS_BY_DOXORUBICIN_DN | 1775 | Genes down-regulated in ME-A cells (breast cancer) undergoing apoptosis in response to doxorubicin [PubChem=31703]. | 73 | 4.33E-17 |
| NAKAYA_PBMC_FLUMIST_AGE_18_50YO_7DY_UP | 1575 | Genes up-regulated in peripheral blood mononuclear cell 7d vs 0d in adults (18-50) after exposure to FluMist , time point 7D. Comment: Supplementary Table 1b: All the differentially expressed genes identified in PBMCs of TIV vaccinees. | 67 | 3.04E-16 |
| BUSSLINGER_GASTRIC_IMMUNE_CELLS | 1492 | - | 65 | 3.04E-16 |
| GOCC_NUCLEAR_PROTEIN_CONTAINING_COMPLEX | 1243 | A stable assembly of two or more macromolecules, i.e. proteins, nucleic acids, carbohydrates or lipids, in which at least one component is a protein and the constituent parts function together in the nucleus. [GOC:pg] | 58 | 1.36E-15 |
| ACEVEDO_LIVER_CANCER_UP | 972 | Genes up-regulated in hepatocellular carcinoma (HCC) compared to normal liver samples. | 49 | 3.96E-14 |

**Table S17.** Top ten most significantly enriched genesets of non-computational collections (C1, C2, C3, C5, C6, C7, C8, H) for the top 500 genes of miRNA-gene pairs with most significant corrected p-values in IDC setting TNBN.

| Geneset Name | # Genes in Geneset | Description | # Genes in Overlap | FDR q-value |
|---|---|---|---|---|
| SMID_BREAST_CANCER_BASAL_DN | 699 | Genes down-regulated in basal subtype of breast cancer samles. | 76 | 3.98E-45 |
| TRAVAGLINI_LUNG_PROXIMAL_CILIATED_CELL | 1770 | - | 89 | 3.62E-27 |
| FARMER_BREAST_CANCER_BASAL_VS_LULMINAL | 329 | Genes which best discriminated between two groups of breast cancer according to the status of ESR1 and AR [GeneID=2099;367]: basal (ESR1- AR-) and luminal (ESR1+ AR+). | 40 | 3.21E-24 |
| CHARAFE_BREAST_CANCER_LUMINAL_VS_BASAL_UP | 384 | Genes up-regulated in luminal-like breast cancer cell lines compared to the basal-like ones. | 42 | 7.73E-24 |
| CHARAFE_BREAST_CANCER_LUMINAL_VS_MESENCHYMAL_UP | 454 | Genes up-regulated in luminal-like breast cancer cell lines compared to the mesenchymal-like ones. | 44 | 5.17E-23 |
| DODD_NASOPHARYNGEAL_CARCINOMA_UP | 1813 | Genes up-regulated in nasopharyngeal carcinoma (NPC) compared to the normal tissue. | 81 | 2.05E-21 |
| GRAESSMANN_APOPTOSIS_BY_DOXORUBICIN_DN | 1775 | Genes down-regulated in ME-A cells (breast cancer) undergoing apoptosis in response to doxorubicin [PubChem=31703]. | 77 | 1.85E-19 |
| NFE2L1_TARGET_GENES | 1913 | Genes containing one or more binding sites for UniProt:Q14494 (NFE2L1) in their promoter regions (TSS -1000,+100 bp) as identified by GTRD version 20.06 ChIP-seq harmonization. | 80 | 2.02E-19 |
| VANTVEER_BREAST_CANCER_ESR1_UP | 149 | Up-regulated genes from the optimal set of 550 markers discriminating breast cancer samples by ESR1 [GeneID=2099] expression: ER(+) vs ER(-) tumors. | 26 | 2.53E-19 |
| BRCA2_TARGET_GENES | 1674 | Genes containing one or more binding sites for UniProt:P51587 (BRCA2) in their promoter regions (TSS -1000,+100 bp) as identified by GTRD version 20.06 ChIP-seq harmonization. | 73 | 1.5E-18 |

**Table S18.** Top ten most significantly enriched genesets of non-computational collections (C1, C2, C3, C5, C6, C7, C8, H) for the top 500 genes of miRNA-gene pairs with most significant corrected p-values in IDC setting TBN.

| Geneset Name | # Genes in Geneset | Description | # Genes in Overlap | FDR q-value |
|---|---|---|---|---|
| SMID_BREAST_CANCER_BASAL_DN | 699 | Genes down-regulated in basal subtype of breast cancer samles. | 71 | 7.99E-40 |
| RODRIGUES_THYROID_CARCINOMA_POORLY_DIFFERENTIATED_DN | 802 | Genes down-regulated in poorly differentiated thyroid carcinoma (PDTC) compared to normal thyroid tissue. | 57 | 2.83E-23 |
| GRAESSMANN_APOPTOSIS_BY_DOXORUBICIN_DN | 1775 | Genes down-regulated in ME-A cells (breast cancer) undergoing apoptosis in response to doxorubicin [PubChem=31703]. | 80 | 4.82E-21 |
| CHARAFE_BREAST_CANCER_LUMINAL_VS_BASAL_UP | 384 | Genes up-regulated in luminal-like breast cancer cell lines compared to the basal-like ones. | 36 | 9.58E-18 |
| NAKAYA_PLASMACYTOID_DENDRITIC_CELL_FLUMIST_AGE_18_50YO_7DY_UP | 1215 | Genes up-regulated in plasmacytoid dendritic cell 7d vs 0d in young adults (18-50) after exposure to FluMist , time point 7D | 59 | 2.62E-16 |
| SCGGAAGY_ELK1_02 | 1242 | Genes having at least one occurrence of the highly conserved motif M3 SCGGAAGY in the regions spanning 4 kb centered on their transcription starting sites [-2kb, +2kb]. This matches the ELK1 [GeneSymbol=ELK1] transcription factor binding site V$ELK1_02 (v7.4 TRANSFAC). | 59 | 6.2E-16 |
| ZNF407_TARGET_GENES | 1996 | Genes containing one or more binding sites for UniProt:Q9C0G0 (ZNF407) in their promoter regions (TSS -1000,+100 bp) as identified by GTRD version 20.06 ChIP-seq harmonization. | 76 | 7.42E-16 |
| TGGAAA_NFAT_Q4_01 | 1935 | Genes having at least one occurrence of the highly conserved motif M55 TGGAAA in the regions spanning 4 kb centered on their transcription starting sites [-2kb, +2kb]. This matches the NFAT [GeneSymbol=NFAT], NFATC [GeneSymbol=NFATC] transcription factor binding site V$NFAT_Q4_01 (v7.4 TRANSFAC). | 74 | 1.54E-15 |
| KRIGE_RESPONSE_TO_TOSEDOSTAT_24HR_UP | 780 | Genes up-regulated in HL-60 cells (acute promyelocytic leukemia, APL) after treatment with the aminopeptidase inhibitor tosedostat (CHR-2797) [PubChem=15547703] for 24 h. | 46 | 1.54E-15 |
| ZNF711_TARGET_GENES | 1762 | Genes containing one or more binding sites for UniProt:Q9Y462 (ZNF711) in their promoter regions (TSS -1000,+100 bp) as identified by GTRD version 20.06 ChIP-seq harmonization. | 70 | 1.81E-15 |

**Table S19.** Top ten most significantly enriched genesets of non-computational collections (C1, C2, C3, C5, C6, C7, C8, H) for the top 500 genes of miRNA-gene pairs with most significant corrected p-values in IDC setting ALLT.

| Geneset Name | # Genes in Geneset | Description | # Genes in Overlap | FDR q-value |
|---|---|---|---|---|
| SMID_BREAST_CANCER_BASAL_DN | 699 | Genes down-regulated in basal subtype of breast cancer samles. | 74 | 4.14E-43 |
| CHARAFE_BREAST_CANCER_LUMINAL_VS_MESENCHYMAL_UP | 454 | Genes up-regulated in luminal-like breast cancer cell lines compared to the mesenchymal-like ones. | 61 | 3.24E-41 |
| VANTVEER_BREAST_CANCER_ESR1_UP | 149 | Up-regulated genes from the optimal set of 550 markers discriminating breast cancer samples by ESR1 [GeneID=2099] expression: ER(+) vs ER(-) tumors. | 35 | 3.35E-31 |
| FARMER_BREAST_CANCER_BASAL_VS_LULMINAL | 329 | Genes which best discriminated between two groups of breast cancer according to the status of ESR1 and AR [GeneID=2099;367]: basal (ESR1- AR-) and luminal (ESR1+ AR+). | 44 | 6.09E-29 |
| TRAVAGLINI_LUNG_PROXIMAL_CILIATED_CELL | 1770 | - | 89 | 1.06E-27 |
| CHARAFE_BREAST_CANCER_LUMINAL_VS_BASAL_UP | 384 | Genes up-regulated in luminal-like breast cancer cell lines compared to the basal-like ones. | 45 | 2.63E-27 |
| DODD_NASOPHARYNGEAL_CARCINOMA_UP | 1813 | Genes up-regulated in nasopharyngeal carcinoma (NPC) compared to the normal tissue. | 79 | 2.71E-20 |
| LIM_MAMMARY_STEM_CELL_DN | 416 | Genes consistently down-regulated in mammary stem cells both in mouse and human species. | 37 | 6.88E-18 |
| FXR1_TARGET_GENES | 1217 | Genes containing one or more binding sites for UniProt:P51114 (FXR1) in their promoter regions (TSS -1000,+100 bp) as identified by GTRD version 20.06 ChIP-seq harmonization. | 58 | 6.13E-16 |
| RODRIGUES_THYROID_CARCINOMA_POORLY_DIFFERENTIATED_DN | 802 | Genes down-regulated in poorly differentiated thyroid carcinoma (PDTC) compared to normal thyroid tissue. | 47 | 6.13E-16 |

**Table S20.** Top ten most significantly enriched genesets of oncogenic collection (C6) for the top 500 genes of miRNA-gene pairs with most significant corrected p-values in LGG setting TNBN.

| Geneset Name | # Genes in Geneset | Description | # Genes in Overlap | FDR q-value |
|---|---|---|---|---|
| KRAS.KIDNEY_UP.V1_UP | 141 | Genes up-regulated in epithelial kidney cancer cell lines over-expressing an oncogenic form of KRAS [Gene ID=3845] gene. | 15 | 2.77E-8 |
| TBK1.DF_DN | 286 | Genes down-regulated in epithelial lung cancer cell lines upon over-expression of an oncogenic form of KRAS [Gene ID=3845] gene and knockdown of TBK1 [Gene ID=29110] gene by RNAi. | 17 | 6.24E-6 |
| TBK1.DN.48HRS_UP | 50 | Genes up-regulated in epithelial lung cancer cell lines upon over-expression of an oncogenic form of KRAS [Gene ID=3845] gene and knockdown of TBK1 [Gene ID=29110] gene by RNAi. | 6 | 1.34E-3 |
| CAMP_UP.V1_UP | 200 | Genes up-regulated in primary thyrocyte cultures in response to cAMP signaling pathway activation by thyrotropin (TSH). | 11 | 1.34E-3 |
| MTOR_UP.V1_DN | 182 | Genes down-regulated by everolimus [PubChem = 6442177] in prostate tissue. | 10 | 2.48E-3 |
| KRAS.600_UP.V1_UP | 278 | Genes up-regulated in four lineages of epithelial cell lines over-expressing an oncogenic form of KRAS [Gene ID=3845] gene. | 12 | 4.06E-3 |
| KRAS.300_UP.V1_UP | 142 | Genes up-regulated in four lineages of epithelial cell lines over-expressing an oncogenic form of KRAS [Gene ID=3845] gene. | 8 | 7.95E-3 |
| RB_P130_DN.V1_UP | 130 | Genes up-regulated in primary keratinocytes from RB1 and RBL2 [Gene ID=5925, 5934] skin specific knockout mice. | 7 | 1.94E-2 |
| SIRNA_EIF4GI_UP | 95 | Genes up-regulated in MCF10A cells vs knockdown of EIF4G1 [Gene ID=1981] gene by RNAi. | 6 | 1.94E-2 |
| AKT_UP.V1_DN | 187 | Genes down-regulated in mouse prostate by transgenic expression of human AKT1 gene [Gene ID=207] vs controls. | 8 | 3.07E-2 |

**Table S21.** Top ten most significantly enriched genesets of oncogenic collection (C6) for the top 500 genes of miRNA-gene pairs with most significant corrected p-values in LGG setting TBN.

| Geneset Name | # Genes in Geneset | Description | # Genes in Overlap | FDR q-value |
|---|---|---|---|---|
| CAMP_UP.V1_UP | 200 | Genes up-regulated in primary thyrocyte cultures in response to cAMP signaling pathway activation by thyrotropin (TSH). | 16 | 4.55E-7 |
| LTE2_UP.V1_DN | 195 | Genes down-regulated in MCF-7 cells (breast cancer) positive for ESR1 [Gene ID=2099] MCF-7 cells (breast cancer) and long-term adapted for estrogen-independent growth. | 12 | 3E-4 |
| CAMP_UP.V1_DN | 199 | Genes down-regulated in primary thyrocyte cultures in response to cAMP signaling pathway activation by thyrotropin (TSH). | 12 | 3E-4 |
| TBK1.DF_DN | 286 | Genes down-regulated in epithelial lung cancer cell lines upon over-expression of an oncogenic form of KRAS [Gene ID=3845] gene and knockdown of TBK1 [Gene ID=29110] gene by RNAi. | 14 | 4.17E-4 |
| KRAS.KIDNEY_UP.V1_UP | 141 | Genes up-regulated in epithelial kidney cancer cell lines over-expressing an oncogenic form of KRAS [Gene ID=3845] gene. | 9 | 1.55E-3 |
| NFE2L2.V2 | 469 | Genes down-regulated in MEF cells (embryonic fibroblasts) after knockout of NFE2L2 [Gene ID=4780] gene. | 17 | 1.55E-3 |
| RB_DN.V1_UP | 116 | Genes up-regulated in primary keratinocytes from RB1 [Gene ID=5925] skin specific knockout mice. | 8 | 1.94E-3 |
| RAF_UP.V1_DN | 193 | Genes down-regulated in MCF-7 cells (breast cancer) positive for ESR1 [Gene ID=2099] MCF-7 cells (breast cancer) stably over-expressing constitutively active RAF1 [Gene ID=5894] gene. | 10 | 2.48E-3 |
| BMI1_DN_MEL18_DN.V1_UP | 145 | Genes up-regulated in DAOY cells (medulloblastoma) upon knockdown of BMI1 and PCGF2 [Gene ID=648, 7703] genes by RNAi. | 8 | 7.02E-3 |
| E2F1_UP.V1_DN | 187 | Genes down-regulated in mouse fibroblasts over-expressing E2F1 [Gene ID=1869] gene. | 9 | 7.5E-3 |

**Table S22.** Top ten most significantly enriched genesets of oncogenic collection (C6) for the top 500 genes of miRNA-gene pairs with most significant corrected p-values in LGG setting ALLT.

| Geneset Name | # Genes in Geneset | Description | # Genes in Overlap | FDR q-value |
|---|---|---|---|---|
| CAMP_UP.V1_DN | 199 | Genes down-regulated in primary thyrocyte cultures in response to cAMP signaling pathway activation by thyrotropin (TSH). | 11 | 2.53E-3 |
| CAMP_UP.V1_UP | 200 | Genes up-regulated in primary thyrocyte cultures in response to cAMP signaling pathway activation by thyrotropin (TSH). | 11 | 2.53E-3 |
| KRAS.KIDNEY_UP.V1_UP | 141 | Genes up-regulated in epithelial kidney cancer cell lines over-expressing an oncogenic form of KRAS [Gene ID=3845] gene. | 9 | 2.87E-3 |
| TBK1.DF_DN | 286 | Genes down-regulated in epithelial lung cancer cell lines upon over-expression of an oncogenic form of KRAS [Gene ID=3845] gene and knockdown of TBK1 [Gene ID=29110] gene by RNAi. | 12 | 7.51E-3 |
| RB_P130_DN.V1_DN | 136 | Genes down-regulated in primary keratinocytes from RB1 and RBL2 [Gene ID=5925, 5934] skin specific knockout mice. | 8 | 7.97E-3 |
| MTOR_UP.V1_DN | 182 | Genes down-regulated by everolimus [PubChem = 6442177] in prostate tissue. | 9 | 9.84E-3 |
| E2F1_UP.V1_DN | 187 | Genes down-regulated in mouse fibroblasts over-expressing E2F1 [Gene ID=1869] gene. | 9 | 9.84E-3 |
| TGFB_UP.V1_UP | 189 | Genes up-regulated in a panel of epithelial cell lines by TGFB1 [Gene ID=7040]. | 9 | 9.84E-3 |
| SIRNA_EIF4GI_UP | 95 | Genes up-regulated in MCF10A cells vs knockdown of EIF4G1 [Gene ID=1981] gene by RNAi. | 6 | 1.88E-2 |
| EIF4E_UP | 100 | Genes up-regulated in HMEC cells (primary mammary epithelium) upon over-expression of EIF4E [Gene ID=1977] gene. | 6 | 2.21E-2 |

**Table S23.** Top ten most significantly enriched genesets of oncogenic collection (C6) for the top 500 genes of miRNA-gene pairs with most significant corrected p-values in LIHC setting TNBN.

| Geneset Name | # Genes in Geneset | Description | # Genes in Overlap | FDR q-value |
|---|---|---|---|---|
| CYCLIN_D1_KE_.V1_UP | 188 | Genes up-regulated in MCF-7 cells (breast cancer) over-expressing a mutant K112E form of CCND1 [Gene ID=595] gene. | 15 | 1.53E-6 |
| SIRNA_EIF4GI_UP | 95 | Genes up-regulated in MCF10A cells vs knockdown of EIF4G1 [Gene ID=1981] gene by RNAi. | 8 | 1.63E-3 |
| NFE2L2.V2 | 469 | Genes down-regulated in MEF cells (embryonic fibroblasts) after knockout of NFE2L2 [Gene ID=4780] gene. | 17 | 3.18E-3 |
| HOXA9_DN.V1_DN | 190 | Genes down-regulated in MOLM-14 cells (AML) with knockdown of HOXA9 [Gene ID=3205] gene by RNAi vs controls. | 10 | 4.43E-3 |
| RB_P107_DN.V1_UP | 130 | Genes up-regulated in primary keratinocytes from RB1 and RBL1 [Gene ID=5925, 5933] skin specific knockout mice. | 8 | 5.88E-3 |
| EIF4E_UP | 100 | Genes up-regulated in HMEC cells (primary mammary epithelium) upon over-expression of EIF4E [Gene ID=1977] gene. | 7 | 5.88E-3 |
| CYCLIN_D1_UP.V1_UP | 187 | Genes up-regulated in MCF-7 cells (breast cancer) over-expressing CCND1 [Gene ID=595] gene. | 9 | 9.49E-3 |
| E2F1_UP.V1_UP | 188 | Genes up-regulated in mouse fibroblasts over-expressing E2F1 [Gene ID=1869] gene. | 9 | 9.49E-3 |
| CYCLIN_D1_UP.V1_DN | 190 | Genes down-regulated in MCF-7 cells (breast cancer) over-expressing CCND1 [Gene ID=595] gene. | 9 | 9.49E-3 |
| MTOR_UP.N4.V1_UP | 196 | Genes up-regulated in CEM-C1 cells (T-CLL) in comparison of control vs rapamycin (sirolimus) [PubChem=6610346], an mTOR pathway inhibitor. | 9 | 1.07E-2 |

**Table S24.** Top ten most significantly enriched genesets of oncogenic collection (C6) for the top 500 genes of miRNA-gene pairs with most significant corrected p-values in LIHC setting TBN.

| Geneset Name | # Genes in Geneset | Description | # Genes in Overlap | FDR q-value |
|---|---|---|---|---|
| ATF2_UP.V1_DN | 185 | Genes down-regulated in myometrial cells over-expressing ATF2 [Gene ID=1386] gene. | 15 | 6.92E-7 |
| ATF2_S_UP.V1_DN | 187 | Genes down-regulated in myometrial cells over-expressing a shortened splice form of ATF2 [Gene ID=1386] gene. | 15 | 6.92E-7 |
| MTOR_UP.N4.V1_UP | 196 | Genes up-regulated in CEM-C1 cells (T-CLL) in comparison of control vs rapamycin (sirolimus) [PubChem=6610346], an mTOR pathway inhibitor. | 15 | 8.72E-7 |
| RPS14_DN.V1_DN | 186 | Genes down-regulated in CD34+ hematopoietic progenitor cells after knockdown of RPS14 [Gene ID=6208] by RNAi. | 14 | 2.24E-6 |
| CYCLIN_D1_KE_.V1_UP | 188 | Genes up-regulated in MCF-7 cells (breast cancer) over-expressing a mutant K112E form of CCND1 [Gene ID=595] gene. | 14 | 2.24E-6 |
| RAPA_EARLY_UP.V1_DN | 186 | Genes down-regulated in BJAB (lymphoma) cells by rapamycin (sirolimus) [PubChem = 6610346]. | 13 | 1.11E-5 |
| LTE2_UP.V1_UP | 188 | Genes up-regulated in MCF-7 cells (breast cancer) positive for ESR1 [Gene ID=2099] MCF-7 cells (breast cancer) and long-term adapted for estrogen-independent growth. | 13 | 1.11E-5 |
| RAF_UP.V1_UP | 193 | Genes up-regulated in MCF-7 cells (breast cancer) positive for ESR1 [Gene ID=2099] MCF-7 cells (breast cancer) stably over-expressing constitutively active RAF1 [Gene ID=5894] gene. | 13 | 1.31E-5 |
| CYCLIN_D1_UP.V1_UP | 187 | Genes up-regulated in MCF-7 cells (breast cancer) over-expressing CCND1 [Gene ID=595] gene. | 11 | 3.12E-4 |
| SIRNA_EIF4GI_UP | 95 | Genes up-regulated in MCF10A cells vs knockdown of EIF4G1 [Gene ID=1981] gene by RNAi. | 8 | 3.12E-4 |

**Table S25.** Top ten most significantly enriched genesets of oncogenic collection (C6) for the top 500 genes of miRNA-gene pairs with most significant corrected p-values in LIHC setting ALLT.

| Geneset Name | # Genes in Geneset | Description | # Genes in Overlap | FDR q-value |
|---|---|---|---|---|
| CYCLIN_D1_KE_.V1_UP | 188 | Genes up-regulated in MCF-7 cells (breast cancer) over-expressing a mutant K112E form of CCND1 [Gene ID=595] gene. | 14 | 1.01E-5 |
| MYC_UP.V1_UP | 181 | Genes up-regulated in primary epithelial breast cancer cell culture over-expressing MYC [Gene ID=4609] gene. | 13 | 2.3E-5 |
| E2F1_UP.V1_UP | 188 | Genes up-regulated in mouse fibroblasts over-expressing E2F1 [Gene ID=1869] gene. | 12 | 1.54E-4 |
| SIRNA_EIF4GI_UP | 95 | Genes up-regulated in MCF10A cells vs knockdown of EIF4G1 [Gene ID=1981] gene by RNAi. | 8 | 7.58E-4 |
| GCNP_SHH_UP_EARLY.V1_UP | 174 | Genes up-regulated in granule cell neuron precursors (GCNPs) after stimulation with Shh for 3h. | 9 | 8.32E-3 |
| MTOR_UP.N4.V1_UP | 196 | Genes up-regulated in CEM-C1 cells (T-CLL) in comparison of control vs rapamycin (sirolimus) [PubChem=6610346], an mTOR pathway inhibitor. | 9 | 1.66E-2 |
| RB_P107_DN.V1_UP | 130 | Genes up-regulated in primary keratinocytes from RB1 and RBL1 [Gene ID=5925, 5933] skin specific knockout mice. | 7 | 2.32E-2 |
| ESC_J1_UP_LATE.V1_UP | 185 | Genes up-regulated during late stages of differentiation of embryoid bodies from J1 embryonic stem cells. | 8 | 3.04E-2 |
| LTE2_UP.V1_UP | 188 | Genes up-regulated in MCF-7 cells (breast cancer) positive for ESR1 [Gene ID=2099] MCF-7 cells (breast cancer) and long-term adapted for estrogen-independent growth. | 8 | 3.04E-2 |
| HOXA9_DN.V1_DN | 190 | Genes down-regulated in MOLM-14 cells (AML) with knockdown of HOXA9 [Gene ID=3205] gene by RNAi vs controls. | 8 | 3.04E-2 |

**Table S26.** Top ten most significantly enriched genesets of oncogenic collection (C6) for the top 500 genes of miRNA-gene pairs with most significant corrected p-values in KIRC setting TNBN.

| Geneset Name | # Genes in Geneset | Description | # Genes in Overlap | FDR q-value |
|---|---|---|---|---|
| VEGF_A_UP.V1_UP | 195 | Genes up-regulated in HUVEC cells (endothelium) by treatment with VEGFA [Gene ID=7422]. | 17 | 3.8E-8 |
| ERBB2_UP.V1_UP | 190 | Genes up-regulated in MCF-7 cells (breast cancer) positive for ESR1 [Gene ID=2099] and engineered to express ligand-activatable ERBB2 [Gene ID=2064]. | 15 | 8.59E-7 |
| STK33_UP | 286 | Genes up-regulated in NOMO-1 and SKM-1 cells (AML) after knockdown of STK33 [Gene ID=65975] by RNAi. | 17 | 3.41E-6 |
| RPS14_DN.V1_UP | 191 | Genes up-regulated in CD34+ hematopoietic progenitor cells after knockdown of RPS14 [Gene ID=6208] by RNAi. | 14 | 3.41E-6 |
| BMI1_DN_MEL18_DN.V1_UP | 145 | Genes up-regulated in DAOY cells (medulloblastoma) upon knockdown of BMI1 and PCGF2 [Gene ID=648, 7703] genes by RNAi. | 12 | 6.27E-6 |
| TBK1.DF_DN | 286 | Genes down-regulated in epithelial lung cancer cell lines upon over-expression of an oncogenic form of KRAS [Gene ID=3845] gene and knockdown of TBK1 [Gene ID=29110] gene by RNAi. | 15 | 5.3E-5 |
| STK33_NOMO_UP | 290 | Genes up-regulated in NOMO-1 cells (AML) after knockdown of STK33 [Gene ID=65975] by RNAi. | 15 | 5.3E-5 |
| ESC_J1_UP_LATE.V1_UP | 185 | Genes up-regulated during late stages of differentiation of embryoid bodies from J1 embryonic stem cells. | 12 | 5.3E-5 |
| CYCLIN_D1_KE_.V1_DN | 191 | Genes down-regulated in MCF-7 cells (breast cancer) over-expressing a mutant K112E form of CCND1 [Gene ID=595] gene. | 12 | 5.91E-5 |
| KRAS.DF.V1_UP | 191 | Genes up-regulated in epithelial lung cancer cell lines over-expressing an oncogenic form of KRAS [Gene ID=3845] gene. | 12 | 5.91E-5 |

**Table S27.** Top ten most significantly enriched genesets of oncogenic collection (C6) for the top 500 genes of miRNA-gene pairs with most significant corrected p-values in KIRC setting TBN.

| Geneset Name | # Genes in Geneset | Description | # Genes in Overlap | FDR q-value |
|---|---|---|---|---|
| LTE2_UP.V1_UP | 188 | Genes up-regulated in MCF-7 cells (breast cancer) positive for ESR1 [Gene ID=2099] MCF-7 cells (breast cancer) and long-term adapted for estrogen-independent growth. | 15 | 1.49E-6 |
| RAF_UP.V1_DN | 193 | Genes down-regulated in MCF-7 cells (breast cancer) positive for ESR1 [Gene ID=2099] MCF-7 cells (breast cancer) stably over-expressing constitutively active RAF1 [Gene ID=5894] gene. | 13 | 5.26E-5 |
| CAHOY_ASTROCYTIC | 100 | Genes up-regulated in astrocytes. | 9 | 1.84E-4 |
| ESC_V6.5_UP_LATE.V1_UP | 184 | Genes up-regulated during late stages of differentiation of embryoid bodies from V6.5 embryonic stem cells. | 11 | 6.05E-4 |
| EGFR_UP.V1_DN | 196 | Genes down-regulated in MCF-7 cells (breast cancer) positive for ESR1 [Gene ID=2099] and engineered to express ligand-activatable EGFR [Gene ID=1956]. | 11 | 8.36E-4 |
| CAMP_UP.V1_DN | 199 | Genes down-regulated in primary thyrocyte cultures in response to cAMP signaling pathway activation by thyrotropin (TSH). | 11 | 8.36E-4 |
| TBK1.DF_DN | 286 | Genes down-regulated in epithelial lung cancer cell lines upon over-expression of an oncogenic form of KRAS [Gene ID=3845] gene and knockdown of TBK1 [Gene ID=29110] gene by RNAi. | 13 | 1.07E-3 |
| ESC_V6.5_UP_LATE.V1_DN | 180 | Genes down-regulated during late stages of differentiation of embryoid bodies from V6.5 embryonic stem cells. | 10 | 1.29E-3 |
| JAK2_DN.V1_DN | 146 | Genes down-regulated in HEL cells (erythroleukemia) after knockdown of JAK2 [Gene ID=3717] gene by RNAi. | 9 | 1.29E-3 |
| E2F1_UP.V1_DN | 187 | Genes down-regulated in mouse fibroblasts over-expressing E2F1 [Gene ID=1869] gene. | 10 | 1.53E-3 |

**Table S28.** Top ten most significantly enriched genesets of oncogenic collection (C6) for the top 500 genes of miRNA-gene pairs with most significant corrected p-values in KIRC setting ALLT.

| Geneset Name | # Genes in Geneset | Description | # Genes in Overlap | FDR q-value |
|---|---|---|---|---|
| BMI1_DN.V1_UP | 147 | Genes up-regulated in DAOY cells (medulloblastoma) upon knockdown of BMI1 [Gene ID=648] gene by RNAi. | 13 | 4.25E-6 |
| ERBB2_UP.V1_UP | 190 | Genes up-regulated in MCF-7 cells (breast cancer) positive for ESR1 [Gene ID=2099] and engineered to express ligand-activatable ERBB2 [Gene ID=2064]. | 14 | 5.05E-6 |
| RAF_UP.V1_UP | 193 | Genes up-regulated in MCF-7 cells (breast cancer) positive for ESR1 [Gene ID=2099] MCF-7 cells (breast cancer) stably over-expressing constitutively active RAF1 [Gene ID=5894] gene. | 14 | 5.05E-6 |
| BMI1_DN_MEL18_DN.V1_UP | 145 | Genes up-regulated in DAOY cells (medulloblastoma) upon knockdown of BMI1 and PCGF2 [Gene ID=648, 7703] genes by RNAi. | 12 | 7.67E-6 |
| ESC_V6.5_UP_LATE.V1_UP | 184 | Genes up-regulated during late stages of differentiation of embryoid bodies from V6.5 embryonic stem cells. | 12 | 7.64E-5 |
| LTE2_UP.V1_UP | 188 | Genes up-regulated in MCF-7 cells (breast cancer) positive for ESR1 [Gene ID=2099] MCF-7 cells (breast cancer) and long-term adapted for estrogen-independent growth. | 12 | 7.64E-5 |
| CYCLIN_D1_KE_.V1_DN | 191 | Genes down-regulated in MCF-7 cells (breast cancer) over-expressing a mutant K112E form of CCND1 [Gene ID=595] gene. | 12 | 7.64E-5 |
| EGFR_UP.V1_UP | 192 | Genes up-regulated in MCF-7 cells (breast cancer) positive for ESR1 [Gene ID=2099] and engineered to express ligand-activatable EGFR [Gene ID=1956]. | 12 | 7.64E-5 |
| CRX_DN.V1_UP | 135 | Genes up-regulated in retina cells from CRX [Gene ID=1406] knockout mice. | 10 | 9.99E-5 |
| MTOR_UP.V1_UP | 169 | Genes up-regulated by everolimus [PubChem = 6442177] in prostate tissue. | 11 | 1.06E-4 |

**Table S29.** Top ten most significantly enriched genesets of oncogenic collection (C6) for the top 500 genes of miRNA-gene pairs with most significant corrected p-values in KICH setting TNBN.

| Geneset Name | # Genes in Geneset | Description | # Genes in Overlap | FDR q-value |
|---|---|---|---|---|
| CAMP_UP.V1_DN | 199 | Genes down-regulated in primary thyrocyte cultures in response to cAMP signaling pathway activation by thyrotropin (TSH). | 14 | 2.21E-5 |
| CAMP_UP.V1_UP | 200 | Genes up-regulated in primary thyrocyte cultures in response to cAMP signaling pathway activation by thyrotropin (TSH). | 13 | 7.7E-5 |
| RAF_UP.V1_UP | 193 | Genes up-regulated in MCF-7 cells (breast cancer) positive for ESR1 [Gene ID=2099] MCF-7 cells (breast cancer) stably over-expressing constitutively active RAF1 [Gene ID=5894] gene. | 11 | 1.12E-3 |
| ERBB2_UP.V1_DN | 197 | Genes down-regulated in MCF-7 cells (breast cancer) positive for ESR1 [Gene ID=2099] and engineered to express ligand-activatable ERBB2 [Gene ID=2064]. | 11 | 1.12E-3 |
| GCNP_SHH_UP_EARLY.V1_UP | 174 | Genes up-regulated in granule cell neuron precursors (GCNPs) after stimulation with Shh for 3h. | 10 | 1.64E-3 |
| GCNP_SHH_UP_LATE.V1_UP | 181 | Genes up-regulated in granule cell neuron precursors (GCNPs) after stimulation with Shh for 24h. | 10 | 1.91E-3 |
| ESC_J1_UP_LATE.V1_UP | 185 | Genes up-regulated during late stages of differentiation of embryoid bodies from J1 embryonic stem cells. | 10 | 1.96E-3 |
| ERBB2_UP.V1_UP | 190 | Genes up-regulated in MCF-7 cells (breast cancer) positive for ESR1 [Gene ID=2099] and engineered to express ligand-activatable ERBB2 [Gene ID=2064]. | 10 | 2.14E-3 |
| P53_DN.V1_UP | 194 | Genes up-regulated in NCI-60 panel of cell lines with mutated TP53 [Gene ID=7157]. | 10 | 2.26E-3 |
| SIRNA_EIF4GI_UP | 95 | Genes up-regulated in MCF10A cells vs knockdown of EIF4G1 [Gene ID=1981] gene by RNAi. | 7 | 2.5E-3 |

**Table S30.** Top ten most significantly enriched genesets of oncogenic collection (C6) for the top 500 genes of miRNA-gene pairs with most significant corrected p-values in KICH setting TBN.

| Geneset Name | # Genes in Geneset | Description | # Genes in Overlap | FDR q-value |
|---|---|---|---|---|
| RAF_UP.V1_UP | 193 | Genes up-regulated in MCF-7 cells (breast cancer) positive for ESR1 [Gene ID=2099] MCF-7 cells (breast cancer) stably over-expressing constitutively active RAF1 [Gene ID=5894] gene. | 15 | 1.76E-6 |
| CAMP_UP.V1_UP | 200 | Genes up-regulated in primary thyrocyte cultures in response to cAMP signaling pathway activation by thyrotropin (TSH). | 15 | 1.76E-6 |
| MEK_UP.V1_DN | 194 | Genes down-regulated in MCF-7 cells (breast cancer) positive for ESR1 [Gene ID=2099] MCF-7 cells (breast cancer) stably over-expressing constitutively active MAP2K1 [Gene ID=5604] gene. | 12 | 1.97E-4 |
| EGFR_UP.V1_DN | 196 | Genes down-regulated in MCF-7 cells (breast cancer) positive for ESR1 [Gene ID=2099] and engineered to express ligand-activatable EGFR [Gene ID=1956]. | 12 | 1.97E-4 |
| LEF1_UP.V1_DN | 187 | Genes down-regulated in DLD1 cells (colon carcinoma) over-expressing LEF1 [Gene ID=51176]. | 11 | 5.55E-4 |
| PGF_UP.V1_UP | 190 | Genes up-regulated in HUVEC cells (endothelium) by treatment with PGF [Gene ID=5281]. | 11 | 5.55E-4 |
| CRX_NRL_DN.V1_DN | 127 | Genes down-regulated in retina cells from CRX and NRL [Gene ID=1406, 4901] double knockout mice. | 9 | 5.64E-4 |
| TBK1.DF_DN | 286 | Genes down-regulated in epithelial lung cancer cell lines upon over-expression of an oncogenic form of KRAS [Gene ID=3845] gene and knockdown of TBK1 [Gene ID=29110] gene by RNAi. | 12 | 3.61E-3 |
| NRL_DN.V1_DN | 132 | Genes down-regulated in retina cells from NRL [Gene ID=4901] knockout mice. | 8 | 3.61E-3 |
| STK33_NOMO_UP | 290 | Genes up-regulated in NOMO-1 cells (AML) after knockdown of STK33 [Gene ID=65975] by RNAi. | 12 | 3.61E-3 |

**Table S31.** Top ten most significantly enriched genesets of oncogenic collection (C6) for the top 500 genes of miRNA-gene pairs with most significant corrected p-values in KICH setting ALLT.

| Geneset Name | # Genes in Geneset | Description | # Genes in Overlap | FDR q-value |
|---|---|---|---|---|
| CAMP_UP.V1_UP | 200 | Genes up-regulated in primary thyrocyte cultures in response to cAMP signaling pathway activation by thyrotropin (TSH). | 18 | 6.34E-9 |
| ERBB2_UP.V1_UP | 190 | Genes up-regulated in MCF-7 cells (breast cancer) positive for ESR1 [Gene ID=2099] and engineered to express ligand-activatable ERBB2 [Gene ID=2064]. | 13 | 3.85E-5 |
| MEK_UP.V1_UP | 195 | Genes up-regulated in MCF-7 cells (breast cancer) positive for ESR1 [Gene ID=2099] MCF-7 cells (breast cancer) stably over-expressing constitutively active MAP2K1 [Gene ID=5604] gene. | 13 | 3.85E-5 |
| LEF1_UP.V1_DN | 187 | Genes down-regulated in DLD1 cells (colon carcinoma) over-expressing LEF1 [Gene ID=51176]. | 12 | 1.16E-4 |
| GCNP_SHH_UP_EARLY.V1_UP | 174 | Genes up-regulated in granule cell neuron precursors (GCNPs) after stimulation with Shh for 3h. | 10 | 1.64E-3 |
| GCNP_SHH_UP_LATE.V1_UP | 181 | Genes up-regulated in granule cell neuron precursors (GCNPs) after stimulation with Shh for 24h. | 10 | 1.91E-3 |
| PGF_UP.V1_UP | 190 | Genes up-regulated in HUVEC cells (endothelium) by treatment with PGF [Gene ID=5281]. | 10 | 2.45E-3 |
| KRAS.300_UP.V1_DN | 140 | Genes down-regulated in four lineages of epithelial cell lines over-expressing an oncogenic form of KRAS [Gene ID=3845] gene. | 8 | 6.15E-3 |
| E2F1_UP.V1_UP | 188 | Genes up-regulated in mouse fibroblasts over-expressing E2F1 [Gene ID=1869] gene. | 9 | 8.54E-3 |
| VEGF_A_UP.V1_UP | 195 | Genes up-regulated in HUVEC cells (endothelium) by treatment with VEGFA [Gene ID=7422]. | 9 | 9.79E-3 |

**Table S32.** Top ten most significantly enriched genesets of oncogenic collection (C6) for the top 500 genes of miRNA-gene pairs with most significant corrected p-values in ILC setting TNBN.

| Geneset Name | # Genes in Geneset | Description | # Genes in Overlap | FDR q-value |
|---|---|---|---|---|
| MEK_UP.V1_UP | 195 | Genes up-regulated in MCF-7 cells (breast cancer) positive for ESR1 [Gene ID=2099] MCF-7 cells (breast cancer) stably over-expressing constitutively active MAP2K1 [Gene ID=5604] gene. | 15 | 2.51E-6 |
| BMI1_DN.V1_UP | 147 | Genes up-regulated in DAOY cells (medulloblastoma) upon knockdown of BMI1 [Gene ID=648] gene by RNAi. | 11 | 1.18E-4 |
| E2F1_UP.V1_DN | 187 | Genes down-regulated in mouse fibroblasts over-expressing E2F1 [Gene ID=1869] gene. | 12 | 1.18E-4 |
| RB_P107_DN.V1_DN | 126 | Genes down-regulated in primary keratinocytes from RB1 and RBL1 [Gene ID=5925, 5933] skin specific knockout mice. | 10 | 1.18E-4 |
| MEK_UP.V1_DN | 194 | Genes down-regulated in MCF-7 cells (breast cancer) positive for ESR1 [Gene ID=2099] MCF-7 cells (breast cancer) stably over-expressing constitutively active MAP2K1 [Gene ID=5604] gene. | 12 | 1.18E-4 |
| P53_DN.V1_UP | 194 | Genes up-regulated in NCI-60 panel of cell lines with mutated TP53 [Gene ID=7157]. | 12 | 1.18E-4 |
| GCNP_SHH_UP_LATE.V1_UP | 181 | Genes up-regulated in granule cell neuron precursors (GCNPs) after stimulation with Shh for 24h. | 11 | 3.02E-4 |
| E2F1_UP.V1_UP | 188 | Genes up-regulated in mouse fibroblasts over-expressing E2F1 [Gene ID=1869] gene. | 11 | 3.77E-4 |
| MEL18_DN.V1_UP | 141 | Genes up-regulated in DAOY cells (medulloblastoma) upon knockdown of PCGF2 [Gene ID=7703] gene by RNAi. | 9 | 1E-3 |
| GCNP_SHH_UP_LATE.V1_DN | 178 | Genes down-regulated in granule cell neuron precursors (GCNPs) after stimulation with Shh for 24h. | 10 | 1.03E-3 |

**Table S33.** Top ten most significantly enriched genesets of oncogenic collection (C6) for the top 500 genes of miRNA-gene pairs with most significant corrected p-values in ILC setting TBN.

| Geneset Name | # Genes in Geneset | Description | # Genes in Overlap | FDR q-value |
|---|---|---|---|---|
| TBK1.DF_DN | 286 | Genes down-regulated in epithelial lung cancer cell lines upon over-expression of an oncogenic form of KRAS [Gene ID=3845] gene and knockdown of TBK1 [Gene ID=29110] gene by RNAi. | 14 | 1.38E-3 |
| E2F1_UP.V1_DN | 187 | Genes down-regulated in mouse fibroblasts over-expressing E2F1 [Gene ID=1869] gene. | 11 | 1.38E-3 |
| EIF4E_DN | 100 | Genes down-regulated in HMEC cells (primary mammary epithelium) upon over-expression of EIF4E [Gene ID=1977] gene. | 8 | 1.53E-3 |
| CYCLIN_D1_UP.V1_DN | 190 | Genes down-regulated in MCF-7 cells (breast cancer) over-expressing CCND1 [Gene ID=595] gene. | 10 | 3.9E-3 |
| RAF_UP.V1_DN | 193 | Genes down-regulated in MCF-7 cells (breast cancer) positive for ESR1 [Gene ID=2099] MCF-7 cells (breast cancer) stably over-expressing constitutively active RAF1 [Gene ID=5894] gene. | 10 | 3.9E-3 |
| MTOR_UP.N4.V1_DN | 184 | Genes down-regulated in CEM-C1 cells (T-CLL) in comparison of control vs rapamycin (sirolimus) [PubChem=6610346], an mTOR pathway inhibitor. | 9 | 9.65E-3 |
| TGFB_UP.V1_UP | 189 | Genes up-regulated in a panel of epithelial cell lines by TGFB1 [Gene ID=7040]. | 9 | 9.65E-3 |
| CYCLIN_D1_KE_.V1_DN | 191 | Genes down-regulated in MCF-7 cells (breast cancer) over-expressing a mutant K112E form of CCND1 [Gene ID=595] gene. | 9 | 9.65E-3 |
| PKCA_DN.V1_DN | 154 | Genes down-regulated in small intenstine in PRKCA [Gene ID=5578] knockout mice. | 8 | 9.65E-3 |
| LEF1_UP.V1_UP | 194 | Genes up-regulated in DLD1 cells (colon carcinoma) over-expressing LEF1 [Gene ID=51176]. | 9 | 9.65E-3 |

**Table S34.** Top ten most significantly enriched genesets of oncogenic collection (C6) for the top 500 genes of miRNA-gene pairs with most significant corrected p-values in ILC setting ALLT.

| Geneset Name | # Genes in Geneset | Description | # Genes in Overlap | FDR q-value |
|---|---|---|---|---|
| TBK1.DF_UP | 287 | Genes up-regulated in epithelial lung cancer cell lines upon over-expression of an oncogenic form of KRAS [Gene ID=3845] gene and knockdown of TBK1 [Gene ID=29110] gene by RNAi. | 18 | 1.93E-6 |
| RAF_UP.V1_DN | 193 | Genes down-regulated in MCF-7 cells (breast cancer) positive for ESR1 [Gene ID=2099] MCF-7 cells (breast cancer) stably over-expressing constitutively active RAF1 [Gene ID=5894] gene. | 13 | 4.53E-5 |
| CAMP_UP.V1_DN | 199 | Genes down-regulated in primary thyrocyte cultures in response to cAMP signaling pathway activation by thyrotropin (TSH). | 13 | 4.53E-5 |
| STK33_UP | 286 | Genes up-regulated in NOMO-1 and SKM-1 cells (AML) after knockdown of STK33 [Gene ID=65975] by RNAi. | 12 | 7.23E-3 |
| BCAT_BILD_ET_AL_DN | 46 | Genes down-regulated in primary epithelial breast cancer cell culture over-expressing activated CTNNB1 [Gene ID=1499] gene. | 5 | 7.34E-3 |
| MTOR_UP.N4.V1_UP | 196 | Genes up-regulated in CEM-C1 cells (T-CLL) in comparison of control vs rapamycin (sirolimus) [PubChem=6610346], an mTOR pathway inhibitor. | 9 | 1.66E-2 |
| RB_P107_DN.V1_UP | 130 | Genes up-regulated in primary keratinocytes from RB1 and RBL1 [Gene ID=5925, 5933] skin specific knockout mice. | 7 | 2.32E-2 |
| GCNP_SHH_UP_EARLY.V1_UP | 174 | Genes up-regulated in granule cell neuron precursors (GCNPs) after stimulation with Shh for 3h. | 8 | 2.49E-2 |
| PGF_UP.V1_UP | 190 | Genes up-regulated in HUVEC cells (endothelium) by treatment with PGF [Gene ID=5281]. | 8 | 3.14E-2 |
| HOXA9_DN.V1_UP | 192 | Genes up-regulated in MOLM-14 cells (AML) with knockdown of HOXA9 [Gene ID=3205] gene by RNAi vs controls. | 8 | 3.14E-2 |

**Table S35.** Top ten most significantly enriched genesets of oncogenic collection (C6) for the top 500 genes of miRNA-gene pairs with most significant corrected p-values in IDC setting TNBN.

| Geneset Name | # Genes in Geneset | Description | # Genes in Overlap | FDR q-value |
|---|---|---|---|---|
| HOXA9_DN.V1_UP | 192 | Genes up-regulated in MOLM-14 cells (AML) with knockdown of HOXA9 [Gene ID=3205] gene by RNAi vs controls. | 13 | 5.26E-5 |
| RAF_UP.V1_DN | 193 | Genes down-regulated in MCF-7 cells (breast cancer) positive for ESR1 [Gene ID=2099] MCF-7 cells (breast cancer) stably over-expressing constitutively active RAF1 [Gene ID=5894] gene. | 13 | 5.26E-5 |
| SIRNA_EIF4GI_UP | 95 | Genes up-regulated in MCF10A cells vs knockdown of EIF4G1 [Gene ID=1981] gene by RNAi. | 9 | 1.2E-4 |
| MTOR_UP.N4.V1_DN | 184 | Genes down-regulated in CEM-C1 cells (T-CLL) in comparison of control vs rapamycin (sirolimus) [PubChem=6610346], an mTOR pathway inhibitor. | 11 | 6.05E-4 |
| ERBB2_UP.V1_UP | 190 | Genes up-regulated in MCF-7 cells (breast cancer) positive for ESR1 [Gene ID=2099] and engineered to express ligand-activatable ERBB2 [Gene ID=2064]. | 11 | 6.54E-4 |
| GCNP_SHH_UP_EARLY.V1_DN | 167 | Genes down-regulated in granule cell neuron precursors (GCNPs) after stimulation with Shh for 3h. | 10 | 9.81E-4 |
| TBK1.DF_DN | 286 | Genes down-regulated in epithelial lung cancer cell lines upon over-expression of an oncogenic form of KRAS [Gene ID=3845] gene and knockdown of TBK1 [Gene ID=29110] gene by RNAi. | 13 | 1.07E-3 |
| MTOR_UP.V1_DN | 182 | Genes down-regulated by everolimus [PubChem = 6442177] in prostate tissue. | 10 | 1.52E-3 |
| PGF_UP.V1_UP | 190 | Genes up-regulated in HUVEC cells (endothelium) by treatment with PGF [Gene ID=5281]. | 10 | 1.94E-3 |
| CAMP_UP.V1_UP | 200 | Genes up-regulated in primary thyrocyte cultures in response to cAMP signaling pathway activation by thyrotropin (TSH). | 10 | 2.66E-3 |

**Table S36.** Top ten most significantly enriched genesets of oncogenic collection (C6) for the top 500 genes of miRNA-gene pairs with most significant corrected p-values in IDC setting TBN.

| Geneset Name | # Genes in Geneset | Description | # Genes in Overlap | FDR q-value |
|---|---|---|---|---|
| CAMP_UP.V1_DN | 199 | Genes down-regulated in primary thyrocyte cultures in response to cAMP signaling pathway activation by thyrotropin (TSH). | 12 | 9.01E-4 |
| CAMP_UP.V1_UP | 200 | Genes up-regulated in primary thyrocyte cultures in response to cAMP signaling pathway activation by thyrotropin (TSH). | 11 | 1.87E-3 |
| GCNP_SHH_UP_EARLY.V1_DN | 167 | Genes down-regulated in granule cell neuron precursors (GCNPs) after stimulation with Shh for 3h. | 10 | 1.87E-3 |
| STK33_UP | 286 | Genes up-regulated in NOMO-1 and SKM-1 cells (AML) after knockdown of STK33 [Gene ID=65975] by RNAi. | 13 | 1.87E-3 |
| ATF2_UP.V1_DN | 185 | Genes down-regulated in myometrial cells over-expressing ATF2 [Gene ID=1386] gene. | 10 | 2.79E-3 |
| PGF_UP.V1_UP | 190 | Genes up-regulated in HUVEC cells (endothelium) by treatment with PGF [Gene ID=5281]. | 10 | 2.9E-3 |
| ERBB2_UP.V1_DN | 197 | Genes down-regulated in MCF-7 cells (breast cancer) positive for ESR1 [Gene ID=2099] and engineered to express ligand-activatable ERBB2 [Gene ID=2064]. | 10 | 3.02E-3 |
| CSR_EARLY_UP.V1_DN | 126 | Genes down-regulated in early serum response of CRL 2091 cells (foreskin fibroblasts). | 8 | 3.02E-3 |
| TBK1.DF_UP | 287 | Genes up-regulated in epithelial lung cancer cell lines upon over-expression of an oncogenic form of KRAS [Gene ID=3845] gene and knockdown of TBK1 [Gene ID=29110] gene by RNAi. | 12 | 3.57E-3 |
| DCA_UP.V1_DN | 185 | Genes down-regulated in A549 lung carcinoma and M059K glioblastoma cells treated with dichloroacetate [PubChem=6597]. | 9 | 6.12E-3 |

**Table S37.** Top ten most significantly enriched genesets of oncogenic collection (C6) for the top 500 genes of miRNA-gene pairs with most significant corrected p-values in IDC setting ALLT.

| Geneset Name | # Genes in Geneset | Description | # Genes in Overlap | FDR q-value |
|---|---|---|---|---|
| ERBB2_UP.V1_UP | 190 | Genes up-regulated in MCF-7 cells (breast cancer) positive for ESR1 [Gene ID=2099] and engineered to express ligand-activatable ERBB2 [Gene ID=2064]. | 13 | 8.41E-5 |
| MTOR_UP.N4.V1_DN | 184 | Genes down-regulated in CEM-C1 cells (T-CLL) in comparison of control vs rapamycin (sirolimus) [PubChem=6610346], an mTOR pathway inhibitor. | 12 | 1.78E-4 |
| EIF4E_DN | 100 | Genes down-regulated in HMEC cells (primary mammary epithelium) upon over-expression of EIF4E [Gene ID=1977] gene. | 9 | 1.78E-4 |
| MTOR_UP.N4.V1_UP | 196 | Genes up-regulated in CEM-C1 cells (T-CLL) in comparison of control vs rapamycin (sirolimus) [PubChem=6610346], an mTOR pathway inhibitor. | 12 | 1.85E-4 |
| RAF_UP.V1_DN | 193 | Genes down-regulated in MCF-7 cells (breast cancer) positive for ESR1 [Gene ID=2099] MCF-7 cells (breast cancer) stably over-expressing constitutively active RAF1 [Gene ID=5894] gene. | 11 | 7.29E-4 |
| GCNP_SHH_UP_EARLY.V1_DN | 167 | Genes down-regulated in granule cell neuron precursors (GCNPs) after stimulation with Shh for 3h. | 10 | 9.48E-4 |
| STK33_SKM_UP | 279 | Genes up-regulated in SKM-1 cells (AML) after knockdown of STK33 [Gene ID=65975] by RNAi. | 12 | 3.09E-3 |
| SIRNA_EIF4GI_UP | 95 | Genes up-regulated in MCF10A cells vs knockdown of EIF4G1 [Gene ID=1981] gene by RNAi. | 7 | 3.09E-3 |
| TBK1.DF_UP | 287 | Genes up-regulated in epithelial lung cancer cell lines upon over-expression of an oncogenic form of KRAS [Gene ID=3845] gene and knockdown of TBK1 [Gene ID=29110] gene by RNAi. | 12 | 3.45E-3 |
| PDGF_ERK_DN.V1_UP | 145 | Genes up-regulated in SH-SY5Y cells (neuroblastoma) in response to PDGF [Gene ID=] stimulation after pre-treatment with the ERK inhibitors U0126 and PD98059 [PubChem=3006531, 4713]. | 8 | 6.12E-3 |
